# MYC promotes immune-suppression in TNBC via inhibition of IFN signaling

**DOI:** 10.1101/2021.02.24.432659

**Authors:** Dario Zimmerli, Chiara S Brambillasca, Francien Talens, Jinhyuk Bhin, Arkajyoti Bhattacharya, Stacey E.P. Joosten, Ana Moises Da Silva, Max D. Wellenstein, Kelly Kersten, Mart de Boo, Maurits Roorda, Linda Henneman, Roebi de Bruijn, Stefano Annunziato, Eline van der Burg, Anne Paulien Drenth, Catrin Lutz, Marieke van de Ven, Lodewyk Wessels, Karin de Visser, Wilbert Zwart, Rudolf S.N. Fehrmann, Marcel A.T.M. van Vugt, Jos Jonkers

## Abstract

Immune checkpoint inhibitor (ICI) treatment has thus far shown limited efficacy in triple-negative breast cancer (TNBC) patients, presumably due to sparse or unresponsive tumor-infiltrating lymphocytes. We reveal a strong correlation between MYC expression and loss of immune signatures in human TNBC. In mouse models of BRCA1-proficient and -deficient TNBC, MYC overexpression dramatically decreased lymphocyte infiltration in tumors, along with immune signature loss. Likewise, MYC overexpression suppressed inflammatory signaling induced by BRCA1/2 inactivation in human TNBC cell lines. Moreover, MYC overexpression prevented the recruitment and activation of lymphocytes in co-cultures with human and mouse TNBC models. Chromatin immunoprecipitation (ChIP)-sequencing revealed that MYC directly binds promoters of multiple interferon-signaling genes, which were downregulated upon MYC expression. Finally, MYC overexpression suppressed induction of interferon signaling and tumor growth inhibition by a Stimulator of Interferon Genes (STING) agonist. Together, our data reveal that MYC suppresses innate immunity and facilitates immune escape, explaining the poor immunogenicity of MYC-overexpressing TNBCs.

**Statement of Significance:** MYC suppresses recruitment and activation of immune cells in TNBC by repressing the transcription of interferon genes. These findings provide a mechanistic rationale for the association of high MYC expression levels with immune exclusion in human TNBCs, which might underlie the relatively poor response of many TNBCs to ICI.

## Introduction

Breast cancer is among the leading causes of cancer-associated death in women, with a lifetime risk of ∼12.5% (Bray et al., 2018). Triple-negative breast cancers (TNBCs) lack expression of the ER, PR and HER2 receptors. A substantial fraction of TNBCs is defective in DNA repair via homologous recombination (HR) due to genetic or epigenetic inactivation of BRCA1 or other components of the HR pathway (Koboldt et al., 2012). Although TNBC only represents 15–20% of breast carcinomas, distant recurrence and mortality in TNBC are significantly higher when compared to other breast cancer subtypes (Dent et al., 2007). As very few targeted therapeutic options are available for patients with TNBC, radiation and chemotherapy are the current standard-of-care treatments, prompting the need for new and more effective treatments (Lehmann et al., 2016).

Targeting the immune system is increasingly employed as a successful treatment approach for cancer. ICIs have resulted in survival benefits across multiple tumor types, with high mutational load and tumor-infiltrating lymphocytes (TILs) being associated with response (Gong et al., 2018). TNBCs were also reported to have high levels of TILs (Loi et al., 2013), (Ibrahim et al., 2014), which was shown to be predictive of treatment response to conventional chemotherapeutics (Denkert et al., 2015), (Bense et al., 2017). Unfortunately, clinical evaluation of single-agent ICI therapy in patients with TNBC only showed benefit in the minority of cases (Kwa and Adams, 2018), (Polk et al., 2018). The poor efficacy of ICI in TNBC patients is surprising, because TNBCs are characterized by multiple features that are associated with response to ICI, including high levels of TILs and potentially high levels of neo-epitopes due to their frequent DNA repair defects which cause pronounced copy number aberrations and complex rearrangements (Narang et al., 2019).

Recently, inactivation of BRCA1/2 and the ensuing DNA damage was shown to result in accumulation of DNA in the cytosol and subsequent activation of the cyclic GMP-AMP synthase / stimulator of interferon (IFN) genes (cGAS/STING) pathway (Parkes et al., 2017), (Heijink et al., 2019). Originally discovered as an anti-viral pathway responding to non-self DNA in the cytosol (Chen et al., 2016), the cGAS/STING pathway was recently described to also respond to ‘own’ DNA, when outside the nucleus (Mackenzie et al., 2017), (Harding et al., 2017). Interestingly, this innate immune pathway was demonstrated to be required for a robust adaptive anti-tumor immune response (Patel et al., 2017), (Manguso et al., 2017). Apparently, TNBCs, and *BRCA1* mutant tumors in particular, have evolved mechanisms to suppress immune responses induced by neo-antigen expression and cGAS/STING signaling.

Multiple recurring gene alterations in tumor-suppressor genes and oncogenes have been described for TNBC. For instance, mutations in the tumor suppressor *TP53* are commonly found along with *BRCA1* in human TNBC (Annunziato et al., 2019). Also, the transcription factor *MYC,* which resides in the 8q24 locus, is regularly amplified in TNBC and especially in *BRCA1*-mutated TNBC (Annunziato et al., 2019). In line with this, a transcriptional signature associated with *MYC* amplification is correlated with a gene signature of *BRCA1*-deficient breast cancers (Alles et al., 2009). MYC regulates global gene expression and thus promotes proliferation and many other cellular processes (Meyer and Penn, 2008) (Kress et al., 2015). Interestingly, MYC was shown to not only promote transcription of targets, but depending on the associated co-factors can also repress transcription (Wanzel et al., 2003). Notably, recent studies have shown that MYC influences the host tumor microenvironment and immune effectors in liver lung and pancreatic cancer (Kortlever et al., 2017), (Casey et al., 2016), (Sodir et al., 2020), (Muthalagu et al., 2020), suggesting a role for MYC in immune suppression beyond its activity as a mitogen. Interestingly, immunogenomic analysis of human TNBC provided hints that MYC overexpression correlates with low immune infiltration (Xiao et al., 2019). However, functional proof that MYC regulates the immune system in TNBC is still lacking and the potential underlying mechanisms are largely unknown.

Here, we explored whether MYC might directly influence immune evasion in TNBC, using a mouse model that recapitulates key features of TNBC. We show that MYC suppresses STING-IFN signaling in a tumor cell-intrinsic fashion, thereby blunting immune cell invasion in TNBC.

## Results

### MYC expression associates with down-regulation of inflammatory pathways in human breast cancer

In TNBCs, the most frequently aberrated oncogene is *MYC*, which was found to be amplified in 61.27% of all TNBC samples within the The Cancer Genome Atlas (TCGA) database (Figure 1A). *MYC* is also the most commonly amplified oncogene in *BRCA1/2*-mutated breast cancers (Supplementary Fig. S1A, Annunziato et al., 2019). To assess the impact of *MYC* expression on inflammatory signaling in TNBCs, we performed gene set enrichment analysis (GSEA) on RNA sequencing (RNA-seq) data obtained from pre-treatment tumor samples from the TONIC phase II trial (NCT02499367), which evaluates the efficacy of nivolumab after immune induction in TNBC patients (Voorwerk et al., 2019). As expected, *MYC* expression positively correlated with MYC target gene sets and E2F targets (Figure 1B). Interestingly, *MYC* expression negatively correlated with IFN and JAK-STAT signaling, as well as other inflammatory pathways, including ‘IL2-STAT5 signaling’, ‘allograft rejection’ and ‘complement activation’ (Figure 1B, C). This is especially interesting considering the fact that in the TONIC trial, increased TILs and immune scores were associated with higher response rates to ICI treatment (Voorwerk et al., 2019). In an independent TCGA dataset, analysis was performed on gene expression data of breast cancer samples that were stratified for amplification or mutation status of selected oncogenes (Supplementary Fig. S1A, S1B). *MYC* amplification again correlated with suppression of gene sets related to inflammatory signaling (Supplementary Fig. S1C). Of note, also other oncogenes seemed to correlate with reduced IFN and JAK-STAT signaling (Supplementary Fig. S1C). However, most oncogene amplifications co-occurred with *MYC* amplification (Figure S1A) or led to upregulated MYC transcriptional signatures, indicating that these effects may also directly or indirectly reflect effects of MYC.

**Figure 1:**
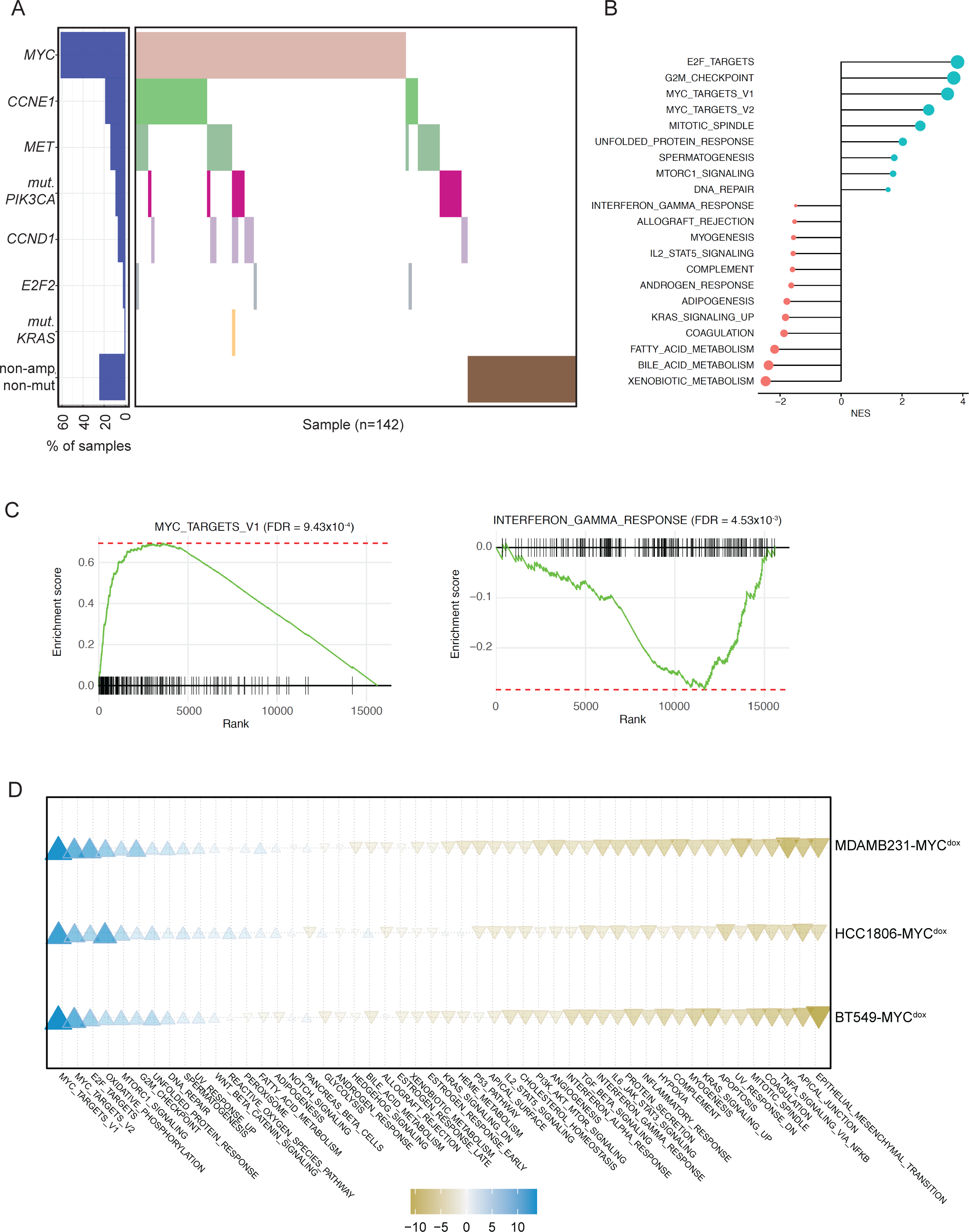
MYC expression associates with down-regulation of inflammatory pathways in human breast cancer. (A) Distribution plot of TNBC samples used for GSEA analysis from TCGA data. In total, 142 samples were included in the analyses. Individual samples are plotted on the x-axis. (B) GSEA for the genes which are positively (cyan) or negatively (pink) correlated with expression of MYC in TONIC trial dataset. The normalized enrichment scores (NES) for the significantly enriched gene sets (FDR<0.05) are presented in the bar plot. MsigDB Hallmark gene sets were used for GSEA analysis. (C) GSEA plots for two significant gene sets, MYC_TARGETS_V1 and INTERFERON_GAMMA_RESPONSE. (D) GSEA of TNBC cell lines BT-549, MB-231 and HCC1806 overexpressing MYC in a dox-inducible manner was performed. Depicted is a bubble plot of the enrichment of a specific gene set between the three tested cell lines upon MYC induction. Increased expression is depicted in blue, while repression is in orange.

To confirm the effects of MYC overexpression on transcriptional re-wiring in an experimental setting, the TNBC cell lines BT-549, MDA-MB-231 and HCC1806 were engineered to overexpress MYC in a doxycycline (dox)-inducible manner. Next, gene expression was analyzed using RNAseq. To investigate the biological processes that are affected by oncogene expression in TNBC cells, GSEA of the gene expression data was performed. Upon MYC overexpression, a strong suppression of IFN signaling pathways was observed, confirming our findings from the human patient samples (Fig. 1D). Taken together, these findings suggest that MYC overexpression suppresses inflammatory signaling, supporting breast cancer to evade detection by the immune system.

### MYC-overexpressing mouse TNBCs display an immune-depleted microenvironment

To explore if MYC regulates immune responses in mammary tumors *in vivo*, we used four genetically engineered mouse models (GEMMs) of BRCA1-proficient and -deficient TBNC with or without engineered MYC overexpression: *WapCre;Trp53^F/F^* (WP), *WapCre;Trp53^F/F^;Col1a1^invCAG-Myc-IRES-Luc/+^* (WP-Myc), *WapCre;Brca1^F/F^;Trp53^F/F^* (WB1P) and *WapCre;Brca1^F/F^;Trp53^F/F^;Col1a1^invCAG-Myc-IRES-Luc/+^* (WB1P-Myc) (Annunziato et al., 2019). GSEA of RNA-seq data of mammary tumors from these four GEMMs showed a clear reduction of immune signatures in the WP-Myc and WB1P-Myc tumors with engineered MYC overexpression, when compared to WP and WB1P control tumors (Figure 2A). Consistently, unsupervised hierarchical clustering of all tumors based on expression of IFN-stimulated genes (ISGs) (Saleiro et al., 2015) resulted in clustering according to MYC status (Supplementary Fig. S2A). Expression of previously published MYC signature genes was consistently increased in our WB1P-MYC versus WB1P models (Bild et al., 2006) confirming the functionality of our MYC-overexpressing mouse models (Supplementary Fig. S2B). WB1P and WP models showed similar change of gene expression profiles MYC expression, indicating the dominant role of MYC in shaping the transcriptional landscape (R=0.67, P<2.2x10^-16^) (Supplementary Fig. S2C). Importantly, immunohistochemical analysis showed a significant reduction of tumor-infiltrating lymphocytes (TILs) in WP-Myc and WB1P-Myc tumors compared to WP and WB1P tumors (Figure 2B, C).

**Figure 2.**
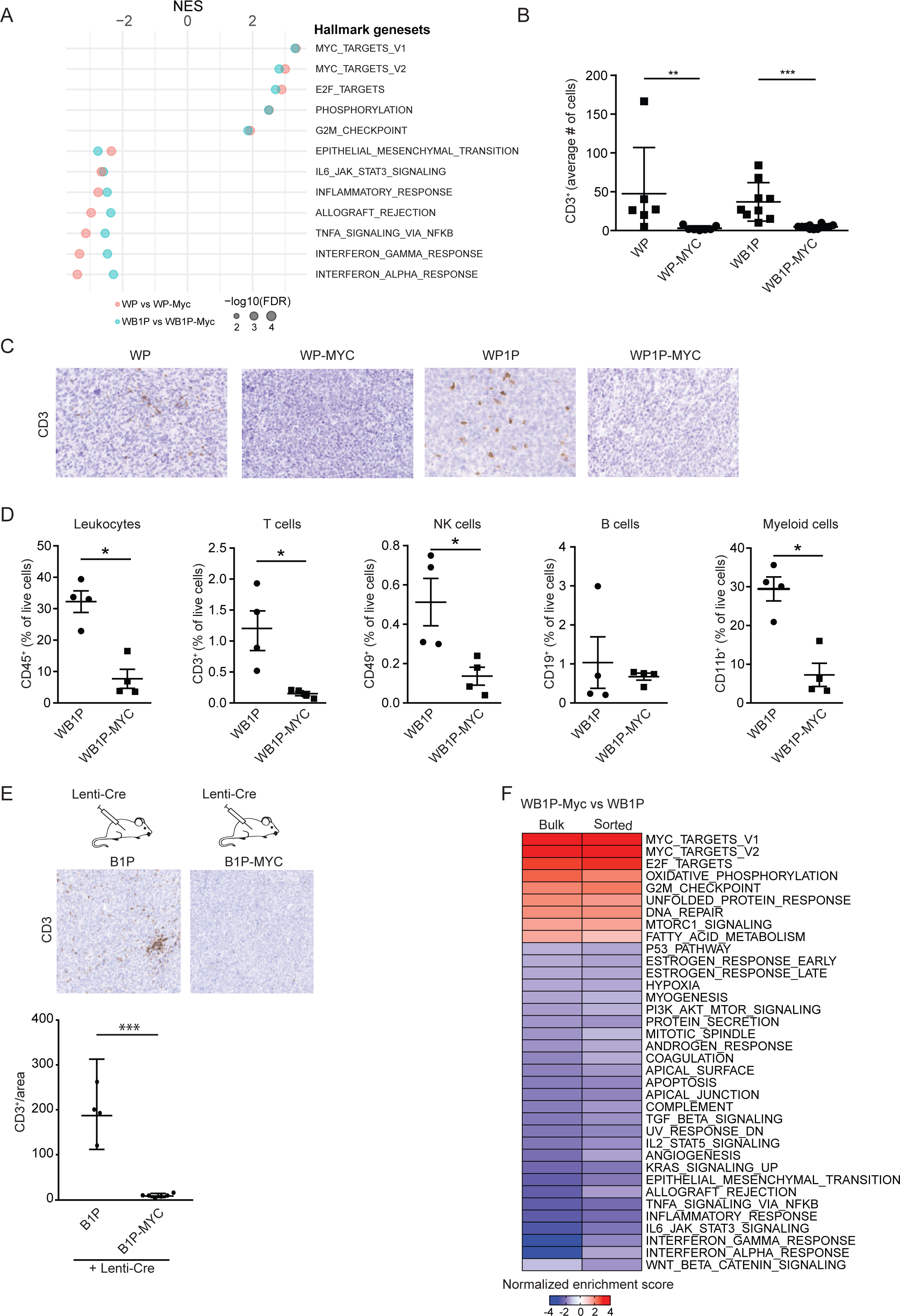
MYC-overexpressing mouse TNBCs display an immune-depleted microenvironment. **(A)** MSigDB Hallmark gene sets significantly represented by WP vs WP-Myc (pink) and WB1P vs WB1P-Myc (cyan) tumors from GSEA analysis. The normalized enrichment scores (NES) for the significantly enriched gene sets (FDR<0.05) are presented in the bar plot. **(B)** Quantification of CD3+ cells in WP, WP-Myc, WB1P and WB1P-Myc tumors (Mann-Whitney test, p=0.0047 and p=0.0001). **(C)** Representative immunostainings for CD3+ cells in WP, WP-Myc, WB1P and WB1P-Myc tumors. **(D)** FACS analysis of T cells (CD3^+^), NK cells (CD49b^+^), B cells (CD19^+^) and myeloid cells (CD11b^+^) in WB1P-Myc tumors in comparison to WB1P tumors (Mann-Whitney test, p<0.05 except for B-cells). **(E)** Top panels: Tumors generated by intraductal lenti-Cre injections in B1P and B1P-Myc mice are analyzed via immunohistochemistry for CD3 expression (brown). Quantifications are shown on the bottom, P values were calculated using two-tailed Student’s t-test, p<0.0001 **(F)** Heatmap for the gene sets represented by the comparison of WB1P-Myc versus WB1P tumor and sorted cancer cells from GSEA analysis. Normalized enrichment scores are plotted for the heatmap.

Since the effect of MYC overexpression on CD3^+^ TILs was most profound in the BRCA1-deficient mammary tumors (Figure 2B), we further focused on the WB1P model. In line with our transcriptomic and histopathologic analysis, flow cytometry analysis of immune cell populations in WB1P-Myc versus WB1P tumors showed a clear loss of CD3^+^ T cells and decreased frequencies of infiltrating CD49b^+^ NK cells and CD11b^+^ myeloid cells in WB1P-Myc tumors (Figure 2D). In contrast, we did not observe a significant difference in CD19^+^ B cell frequencies (Figure 2D). Of note, draining lymph nodes, spleen, and blood showed similar lymphocyte frequencies in WB1P and WB1P-Myc mice, arguing against systemic immune-suppression and pointing towards local dampening of the immune response via paracrine signals from tumor cells (Supplementary Fig. S2D). To further corroborate our findings in WB1P and WB1P-Myc mice, we used somatic engineering (Annunziato et al., 2016) to induce mammary tumors in *Brca1^F/F^;Trp53^F/F^* (B1P) and *Brca1^F/F^;Trp53^F/F^;Col1a1^invCAG-Myc-IRES-Luc/+^* (B1P-Myc) mice via intraductal injection of a Cre-encoding lentivirus. This resulted again in profound TIL depletion in the MYC-overexpressing B1P tumors (Figure 2E).

To test if the reduced immune cell infiltration was not merely due to accelerated tumorigenesis in WB1P-Myc mice compared to WB1P mice, we generated *WapCre;Brca1^F/F^;Trp53^F/F^;Col1a1^invCAG-Met-IRES-Luc/+^* (WB1P-Met) mice with tumor-specific overexpression of MET instead of MYC, leading to a very similar decrease in tumor latency as observed in the WB1P-Myc model (Supplementary Fig. S2E). In contrast to MYC, MET overexpression did not result in immune suppression, as demonstrated by the comparable numbers of TILS in WB1P-Met versus WB1P tumors (Supplementary Fig. S2F). Also clustering based on expression of ISGs resulted in a clear separation between WB1P-Met and WB1P-Myc tumors (Supplementary Fig. S3A). To further examine if the immune cell exclusion in WB1P-Myc tumors was not a generic consequence of tumor-promoting mutations, we tested if loss of an unrelated tumor suppressor, *Pten,* would also lead to decreased lymphocyte infiltration. To this end, intraductal injections were performed in *WapCre;Brca1^F/F^;Trp53^F/F^;Col1a1^invCAG-Cas9-IRES-Luc/+^* (WB1P-Cas9) mice with lentiviruses encoding a *Pten*-targeting sgRNA alone (Lenti-sgPten) or in combination with MYC (Lenti-sgPten-Myc) (Annunziato et al., 2019). TILs were observed in tumors from mice injected with Lenti-sgPten, but not in tumors from mice injected with Lenti-sgPten-Myc, confirming that MYC is selectively responsible for immune cell exclusion (Supplementary Fig. S3B).

### MYC drives immune cell exclusion in a tumor cell-intrinsic manner

To investigate how MYC is linked to an immune-suppressive phenotype, we performed RNA-seq on two different sources of tumor cells. In addition to bulk tumors containing both tumor cells and infiltrating immune cells, we used FACS-sorted E-cadherin-positive (ECAD^+^) tumor cells from WB1P and WB1P-Myc tumors. Consistent with our analysis mentioned above, GSEA showed significant downregulation of immune response pathways in the bulk tumor samples (Figure 2F). Although such transcriptomic changes could be due to the decreased presence of immune cells in bulk tumor samples, these immune pathways were also downregulated in sorted tumor cells, indicating that MYC-associated immune evasion is mediated by a tumor cell-intrinsic mechanism (Figure 2F). In support of this notion, the enriched pathways showed strong correlation (R=0.83) between WB1P-Myc bulk tumors and sorted tumor cells (Supplementary Fig. S3C), further underscoring that MYC suppresses IFN signaling in a tumor cell-intrinsic manner.

To corroborate on our preclinical *in vivo* findings, we used CIBERSORT analysis (Newman et al., 2019) on gene expression data from TCGA to estimate the fractions of different immune cell types in human breast cancer samples. Compared to cancers with neutral copy numbers of *MYC*, breast cancers with amplified *MYC* contained lower fractions of monocytes, M2 macrophages and CD8^+^ T cells, whereas they showed increased fractions of M0/M1 macrophages and regulatory T cells (Supplementary Fig. S3D). A similar pattern was observed within the TNBC subset of breast cancers (Supplementary Fig. S3D). Taken together, our results show that MYC expression drives a dramatic loss of lymphocytic infiltration, as well as other immune cells, in mouse and human breast cancer. Furthermore, we demonstrate a cancer cell-intrinsic role for MYC in suppressing inflammatory pathways.

### MYC overexpression in mammary tumor cells down-regulates IFN-stimulated genes

The main downregulated pathways in the MYC-overexpressing WB1P tumors were ‘IFN signaling’ and ‘JAK/STAT signaling’ (Figure 2A, F), which are both important in inflammatory responses (Villarino et al., 2015). Recent studies showed that loss of BRCA1 leads to accumulation of cytosolic DNA, thereby triggering the cGAS/STING pathway (Ding et al., 2018), (Parkes et al., 2017), (Heijink et al., 2019). We derived organoids from WB1P and WB1P-Myc tumors to investigate whether suppression of IFN signaling in WB1P-Myc tumors is connected to reduced cGAS/STING pathway activation. We first probed RNA-seq profiles from bulk tumors, sorted cells and organoids with a previously reported panel of ISGs induced by cGAS/STING signaling (Mackenzie et al., 2017). The expression of these ISGs clearly separated WB1P-Myc from the WB1P tumors and organoids, showing significant downregulation of the ISGs (Figure 3A).

**Figure 3.**
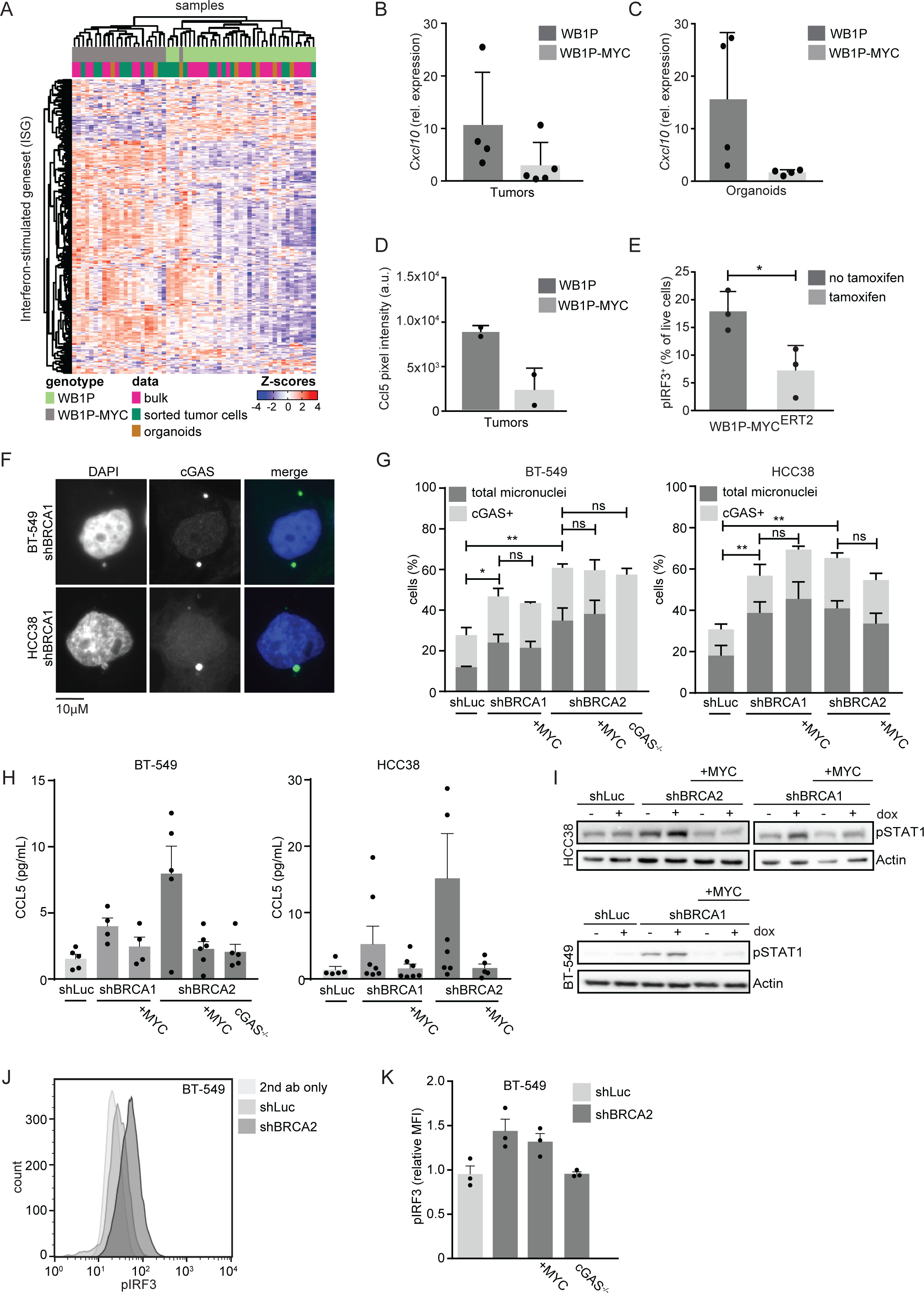
MYC overexpression in mammary tumor cells down-regulates IFN-stimulated genes. (A) Heatmap depicting the previously reported interferon-stimulated genes (Mackenzie et al., 2017) in RNA-seq of bulk tumors, sorted epithelial tumor cells, and organoids. (B) qRT-PCR analysis in mouse tumors for Cxcl10, relative gene expression levels normalized to GAPDH are plotted. (C) qRT-PCR analysis for Cxcl10 in 3 independent mouse tumor organoid lines each plus a line transfected with Myc^ERT2^ with and without Tamoxifen, relative gene expression levels normalized to RPM20 are plotted. (D) Quantification of cytokine array for Ccl5 in WB1P and WB1P-Myc organoids. (n=2). (E) Quantification of pIRF3 by FACS analysis of WB1P-Myc^ERT2^ organoids with and without tamoxifen treatment. (n=3, p= 0.03) P values were calculated using two-tailed Student’s t-test. (F) Representative images of BT-549 and HCC38 cells harboring shBRCA2 and treated with dox for three days. Cells were stained with anti-cGAS and DAPI. (G) Quantification of cGAS-positive micronuclei as described in F. ≥100 Cells were counted per condition. Error bars indicate SEM of at least three independent experiments. P values were calculated using two-tailed Student’s t-test. * P <0.05, **P < 0.01. (H) BT-549 and HCC38 cells with indicated hairpins, depleted for cGAS or overexpressed for MYC were treated with or without dox for 6 days. Supernatants were harvested and secretion of CCL5 was measured with ELISA. Concentrations were normalized to untreated conditions of each cell line. Means are indicated underneath each condition. Error bars indicate SEM of at least 4 independent experiments with two technical replicates each. (I) BT-549 and HCC38 cells with indicated hairpins, depleted for cGAS or overexpressed for MYC were treated with or without dox for 5 days. Phosphorylation status of STAT1 was analyzed by immunoblotting. (J) BT-549 cells with indicated hairpins, were treated with dox for 5 days. Phosphorylation levels of IRF3 were analyzed by flow cytometry. (K) Phosphorylation levels of IRF3 were analyzed by flow cytometry as described in J. Median fluorescence intensities (MFI) were normalized to cells treated without dox. Error bars indicate SEM of at least 3 independent experiments.

Specifically, we observed down-regulation of various STING-pathway-related genes, including *Stat1*, *Stat3*, *Ccl20* and *Irf9* (Supplementary Fig. S7B) (Dunphy et al., 2018), (Platanitis et al., 2019). We also observed downregulation of *Cd74* and *Ciita*, two genes regulated directly by STAT1 and important for the function of the adaptive immune response via MHC class II signaling (Muhlethaler-Mottet et al., 1998) (Supplementary Fig. S4A). Downstream activation of cGAS/STING signaling is associated with the secretion of different chemokines and cytokines, including CCL5 and CXCL10 (Mackenzie et al., 2017), (Motani et al., 2015). In line with this notion, we found that MYC-expressing tumors and organoids show reduced expression of *Cxcl10*, STAT1 and CCL5 (Figure 3B-D, Supplementary Fig. S4B, C). To confirm the direct role of MYC in the downregulation of ISGs, we transduced WB1P organoids with a lentiviral vector encoding a tamoxifen-inducible MYC^ERT2^ fusion protein. Flow cytometry analysis of these organoids showed that MYC activation upon addition of tamoxifen decreased phosphorylation of Interferon Regulatory Factor 3 (pIRF3) (Figure 3E, Supplementary Fig. S4D), a key transcriptional regulator of IFN and STING responses, as well as phosphorylation of Tank binding kinase (pTBK1) (Supplementary Fig. S4E), a central player in the STING signaling pathway. We conclude that MYC overexpression can manipulate IFN signaling by reducing the expression of a broad network of genes in our murine tumor model.

To investigate the effects of MYC overexpression on cGAS/STING signaling in human breast cancer, the TNBC cell lines BT-549 and HCC38 were transduced with dox-inducible short hairpin RNAs (shRNAs) targeting BRCA1 or BRCA2, with or without constitutive overexpression of MYC (Supplementary Fig. S4F), whereas cGAS^-/-^ cells served as controls. Of note, shRNA-mediated depletion of BRCA1 and BRCA2 resulted in decreased cell proliferation and ultimately cell death (Supplementary Fig. S4G-I), which was not rescued by MYC overexpression (Supplementary Fig. S4G-I). In line with previous reports (Mackenzie et al., 2017), (Heijink et al., 2019), BRCA1 and BRCA2 depletion led to increased amounts of cGAS-positive micronuclei (Figure 3F,G). This increase was not suppressed by MYC overexpression (Figure 3G), suggesting that the role of MYC in suppressing IFN signaling acts downstream of the generation of cytoplasmic DNA.

In line with our observations in WB1P and WB1P-Myc mammary tumors, expression of CCL5, IFNβ, IFNγ, CXCL10 and pSTAT1 was induced upon BRCA1/2 depletion, and was suppressed in MYC-overexpressing BT-549 and HCC38 cells (Figure 3H,I, Supplementary Fig. S5A). In contrast, pIRF3 levels were only marginally affected (Figure 3J,K, Supplementary Fig. S5B), suggesting differences between the human TNBC cells and mouse mammary tumor organoids. RNAseq in both BT-549 and HCC38 cell lines with BRCA2-depletion with and without MYC overexpression confirmed the ability of MYC to downregulate inflammatory signaling, including the ‘IL2 STAT5 signaling’ and ‘Interferon gamma response’ pathways (Supplementary Fig. S5C, D).

### MYC status of breast cancer cells regulates lymphocyte trafficking and activation *in vitro* and *in vivo*

To further assess the role of MYC in the inhibition of inflammatory signaling and thereby suppressing immune responses, we turned to human and mouse *in vitro* systems. Using trans-well assays, we measured the migration of isolated human CD8^+^ T cells towards BT-549 and HCC38 cells upon BRCA1 or BRCA2 depletion (Figure 4A). Upon BRCA1/2-depletion for 24h or 48h, we observed increased numbers of CD8^+^ T cells that migrated towards the BT-549 and HCC38 tumor cells, which was blocked upon MYC overexpression or loss of cGAS (Figure 4B). In parallel, we harvested conditioned medium to probe whether secreted factors upon BRCA1/2 depletion conferred an ability to stimulate CD8^+^ T cell proliferation and activation (Supplementary Fig. S5F). While medium from BRCA1- or BRCA2-depleted cells stimulated activation and proliferation, overexpression of MYC had inhibitory effects (Figure 4C). These results suggest that factors secreted by BRCA1- or BRCA2-depleted breast cancer cells increase the migration and activation of T cells via activation of STING signaling. In contrast, MYC overexpression suppresses the migration and activation of CD8^+^ T cells in a tumor cell-intrinsic manner.

**Figure 4.**
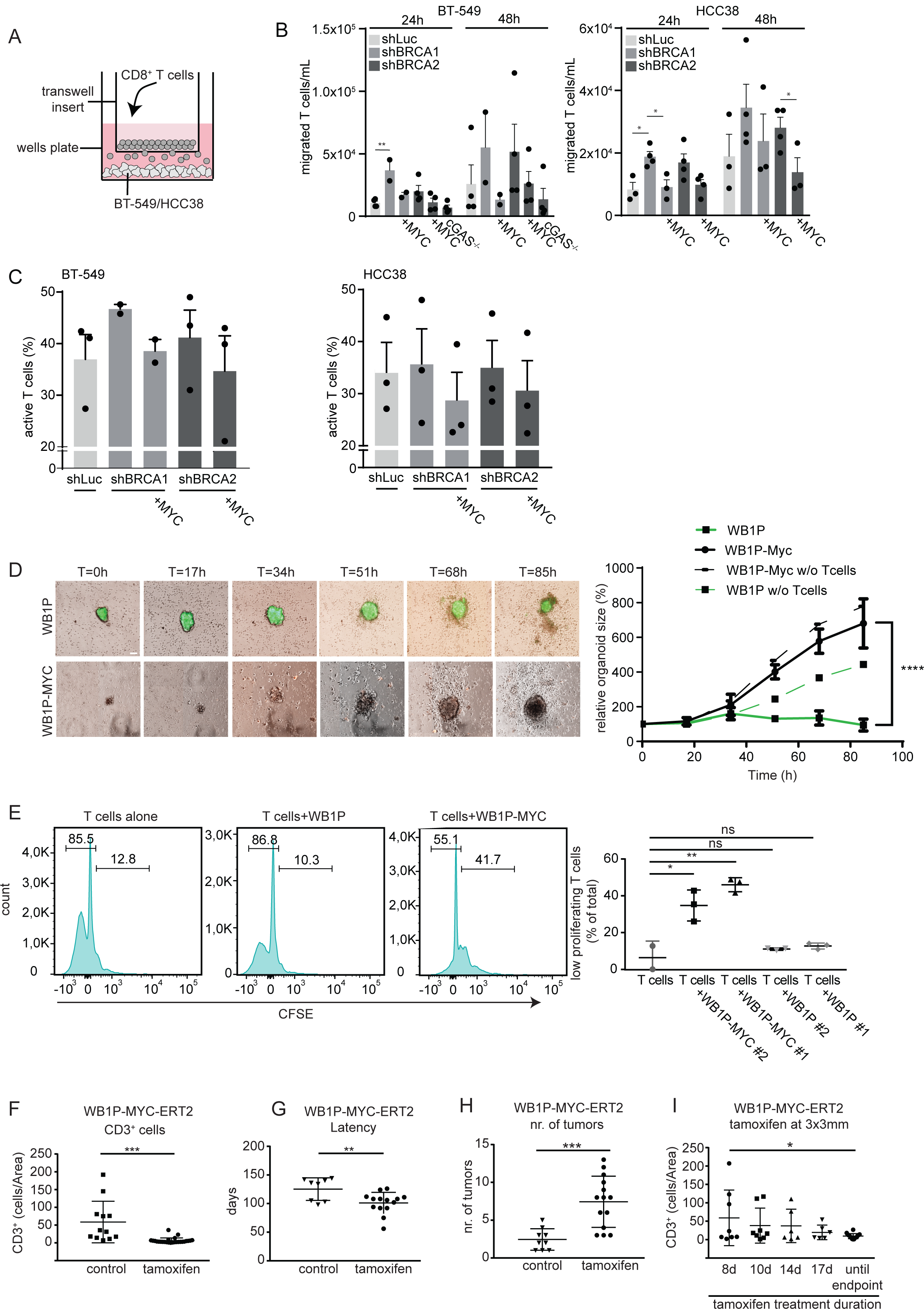
MYC directly regulates lymphocyte trafficking and activation *in vitro* and *in vivo*. (A) Schematic overview of the transwell assay. BT-549 or HCC38 cells were cultured in wells for 5 days with dox treatment. Isolated human CD8^+^ T cells were added in the transwell insert and cultured for 1 or 2 days. The amount of migrated T cells towards the lower compartment were counted. (B) Transwell assays were performed as described in A. The amount of migrated T cells towards the tumor cells was counted after 24 or 48 hours. (C) The percentage of activated CD8^+^ T cells was measured by flow cytometry 5 days after co-culture with harvested medium from BT-549 or HCC38 cells pre-treated with dox. Gating was performed as shown in S5F. Mean percentages are indicated underneath each condition. Error bars indicate SEM of 2 or 3 independent experiments. (D) Live imaging of in vitro co-culture of WB1P (green) and WB1P-Myc organoids (red) with splenocytes. Time after seeding in hours is indicated on top of the respective panels. Quantification of organoid sizes is quantified in the right panel (scale=100μm). (E) FACS analysis of WB1P and WB1P-Myc tumor derived organoids co-cultured with T-cells and stained for CSFE. Halving in fluorescence intensity marks one cell division. Fluorescence intensity per number of cells is plotted. Quantification for percentage of low proliferating Tcells of replicas of 2 different organoid lines each is shown in the right panel. (unpaired t-tests, comparisons: Tcells with Tcells + WB1PMyc line1, p=0.04, Tcells with Tcells + WB1PMyc line2, p=0.01, Tcells with Tcells + WB1P line1, p=0.4, T, Tcells with Tcells+WB1P line2, p=0.29). (F) Counts of CD3+ T-cells of tumors in the Myc^ERT2^ GEMM model with and without tamoxifen chow. (unpaired t-test, p=0.0001). (G) Latency analysis in WB1P-Myc^ERT2^ with and without tamoxifen. (unpaired t-test, p=0.009) (H) Tumor burden in mice with and without tamoxifen administration. (unpaired t-test, p=0.0005). (I) CD3+ cell counts in tumors upon starting administration of tamoxifen when tumors are 3 by 3 mm for the time indicated.(p=0.014, two tailed t-test, comparison Tam for 8 days and until endpoint.)

To confirm that MYC overexpression has similar effects in mouse mammary tumor cells, we performed live-cell imaging in co-cultures of WB1P and WB1P-Myc tumor-derived organoids and syngeneic mouse splenocytes. 7-day time-lapse quantification of organoid size demonstrated clear growth inhibition of WB1P organoids by immune cells, whereas immune cells did not significantly affect WB1P-Myc organoids (Figure 4D). Also, proliferation analysis in these co-cultures demonstrated that WB1P-Myc organoids suppressed IL2-induced T-cell proliferation (Figure 4E). To investigate whether MYC directly controls immune cell tumor infiltration *in vivo*, we assessed the effects of MYC-activation or -inactivation in established BRCA1-deficient mouse mammary tumors *in situ*. To this end, we generated *WapCre;Brca1^F/F^;Trp53^F/F^;Col1a1^invCAG-MycERT2-IRES-Luc/+^* (WB1P-Myc^ERT2^) GEMMs and investigated tumor growth and immune infiltration at different time points after MYC induction via tamoxifen administration. Upon feeding tamoxifen at seven weeks of age until the time of sacrifice, we again observed that MYC induction reduced immune cell infiltration and shortened tumor latency compared to WB1P tumors, similar to findings with the WB1P-Myc model (Figure 4F,G), (Supplementary Fig. S6A). Also, we observed higher numbers of tumors, as expected for MYC-driven tumorigenesis (Figure 4H). Whereas tumor growth rates were not significantly altered upon tamoxifen-induced Myc^ERT2^ translocation in already established tumors, MYC activation resulted in depletion of immune infiltrates (Figure 4I). Specifically, a time-dependent reduction in CD3^+^ T-cells was observed after tamoxifen administration until they resembled the low levels that were observed in WB1P-Myc mice (Figure 4I), (Supplementary Fig. S6A). MYC-induced reduction of TILs was also observed in B1P-Myc^ERT2^ tumors induced by intraductal injection of lentiviral Myc^ERT2^-P2A-Cre as well as in orthotopically transplanted WB1P tumor-derived organoids that were transduced with lentiviral Myc^ERT2^ (Supplementary Fig. S6B-D). Combined, these observations confirmed our previous findings that MYC expression directly hinders immune infiltration in BRCA1-mutant tumors and underscore that the decreased immunogenicity does not result from different tumor latencies.

### MYC controls expression of multiple IFN signaling components in tumors and organoids

To test if MYC directly regulates genes involved in IFN signaling, we performed chromatin immuno-precipitation of MYC, followed by massive parallel sequencing (ChIP-seq) on WB1P and WB1P-Myc organoids as well as tumors. We found 1257 shared peaks between the tumor and organoid ChIPs (Fig. 5A), of which the majority was found in promoter regions (Figure 5B). As expected, analysis of the enriched motif sequences within the ChIP-seq peaks revealed that most peaks were found in the promoter area and contained the Myc binding motif (Supplementary Fig. S7C). MYC binding was significantly enriched in the promoter regions of those genes that were previously found to be up-regulated in the RNAseq of WB1P-Myc bulk tumors, sorted tumor cells and organoids (Figure 5C).

**Figure 5.**
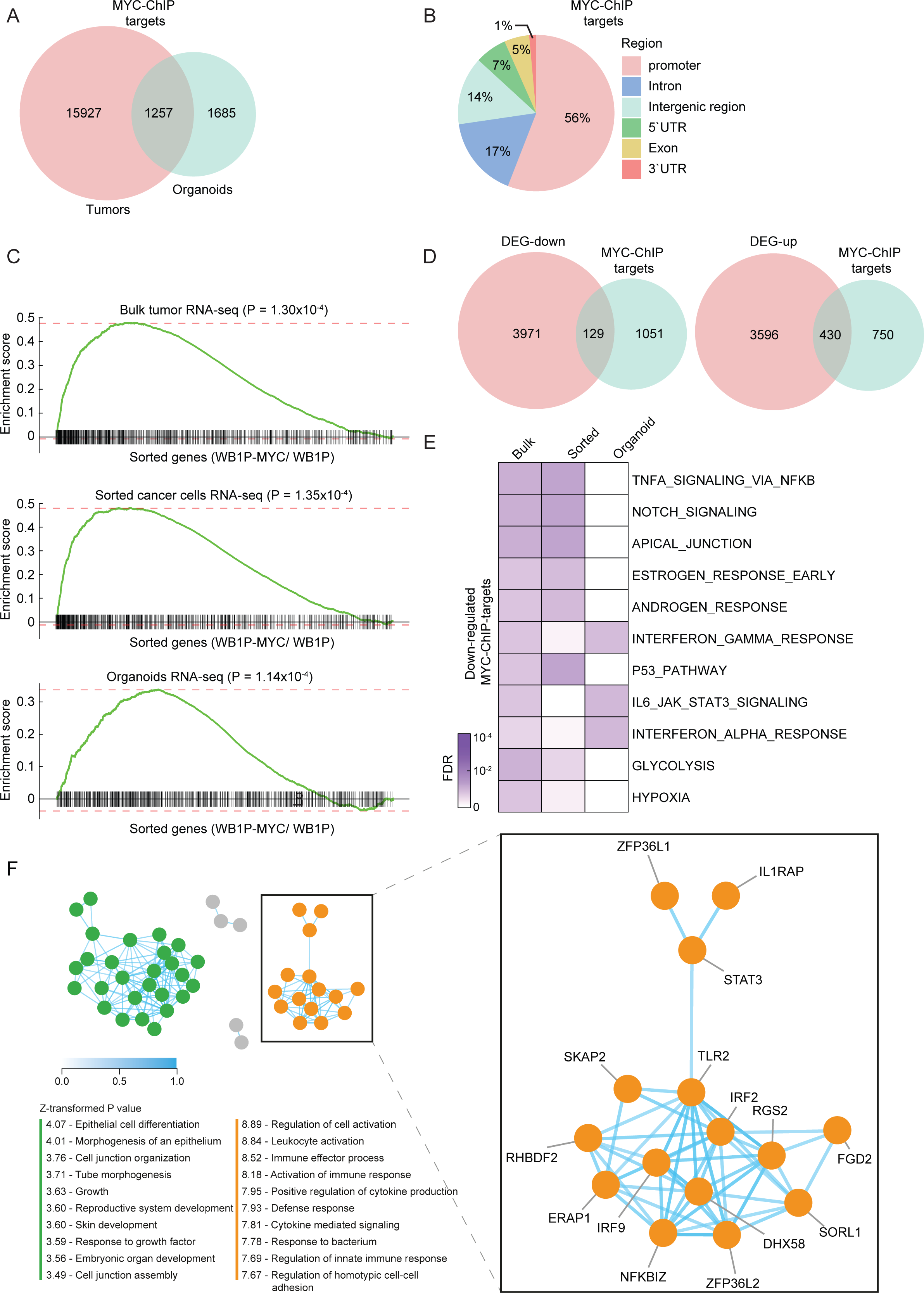
MYC controls expression of multiple IFN signaling components in tumors and organoids. (A) Overlap of the MYC-binding loci obtained from the tumor and organoid ChIP-seq data (B) Genomic distribution of the common Myc-binding loci between tumor and organoid ChIP-seq data. (C) GSEA analysis of Myc targets in each bulk tumor, sorted cancer cells, and organoids RNA-seq data comparing WB1P-Myc to WB1P. Myc targets were defined by the genes nearest to common Myc-binding loci between tumor and organoid ChIP-seq. (D) Overlap between MYC target genes from ChIP-seq and differentially expressed genes (DEGs) in the comparison of WB1P-Myc versus WB1P. For the comparison, MYC targets were obtained from the common MYC-binding loci from tumor and organoid ChIP-seq data, and DEGs were obtained from the union of the genes showing differential expression between WB1P-Myc versus WB1P in bulk tumor, sorted tumor cells, and organoid RNA-seq data (see Methods). (E) Gene sets significantly represented by the down-regulated Myc targets from Fisher’s exact test (FDR<0.1). The down-regulated Myc targets were defined by the genes in Figure 5D (129 down-regulated Myc targets). (F) Left: constructed co-functionality network of genes downregulated by MYC (n=129) retrieved from both MYC-ChIP-seq of WB1P-Myc and RNA-seq data of WB1P and WB1P-Myc tumors and organoids. Right: one of the two resulting clusters in which genes share strong predicted co-functionality (r > 0.5) and show enrichment for immunity pathways (e.g. leukocyte activation, activation of immune response and positive regulation of cytokine secretion).

Next, we intersected the MYC-bound genes identified by ChIP-seq with the genes downregulated upon MYC overexpression in the mouse mammary tumors, organoids and sorted tumor cells (Figure 5D, Supplementary Fig. S7A, S7D). GSEA revealed that genes involved in IFN signaling and inflammation, including ‘JAK STAT3 signaling’ (Figure 5E) were enriched among the 129 genes that were bound by MYC and downregulated upon MYC overexpression. MYC ChIP-seq peaks that occurred in tumors and/or organoids overlapped with 59 IFN signaling pathway-associated genes that were repressed by MYC in the tumors, sorted tumor epithelial cells and/or organoids (Supplementary Fig. S7B). We next constructed a co-functionality network (Bhattacharya et al., 2020) using all 129 MYC-repressed genes (Figure 5D), which revealed a network of immune- and IFN-related genes among the MYC target genes, confirming the role of MYC in suppressing inflammatory signaling (Figure 5F). Conversely, a co-functionality network for MYC-upregulated genes did not show any immunity signatures (Supplementary Fig. S7E). Combined, our results demonstrate that MYC directly controls immune infiltration into tumors via downregulation of a myriad of inflammatory pathway components.

### MYC suppresses anti-cancer efficacy of pharmacological STING activation in WB1P tumors

To further investigate how MYC influences inflammatory signaling, we used the small molecule STING agonist vadimezan (DMXAA) to activate IFN signaling in mice with established WB1P and WB1P-Myc tumors. WB1P and WB1P-Myc organoids were transplanted into mammary fat pads of immunocompetent syngeneic mice, and a single dose of vadimezan was administered to the mice when tumor outgrowths reached a volume of 100 mm^3^. Vadimezan treatment induced transient regression of WB1P tumors for one to two weeks, after which tumor growth resumed (Figure 6A). The same experiment using transplanted WB1P-Myc organoids showed that these tumors were more resistant to STING activation than WB1P tumors, as tumor regression was not observed in this setting. Of note, vademizan-treated WB1P-Myc tumors still showed a clear growth delay in comparison to vehicle-treated tumors (Figure 6B). Similarly, STING-agonist treatment of co-cultures of tumor organoids with splenocytes strongly blocked growth of WB1P organoids as measured by MTT assay, whereas WB1P-Myc organoids were largely unaffected, confirming our *in vivo* experiments (Figure 6C). Together, these findings show the relevance of IFN signaling in suppressing tumor growth in TNBCs and underscore the critical role of MYC in undermining the IFN response via direct inhibition of immunomodulatory factors downstream and in parallel of c-GAS/STING signaling. Our findings also highlight the difficulty of countering the broad immunoinhibitory effects of MYC by therapeutic targeting of a single downstream factor such as c-GAS/STING (Figure 6D).

**Figure 6.**
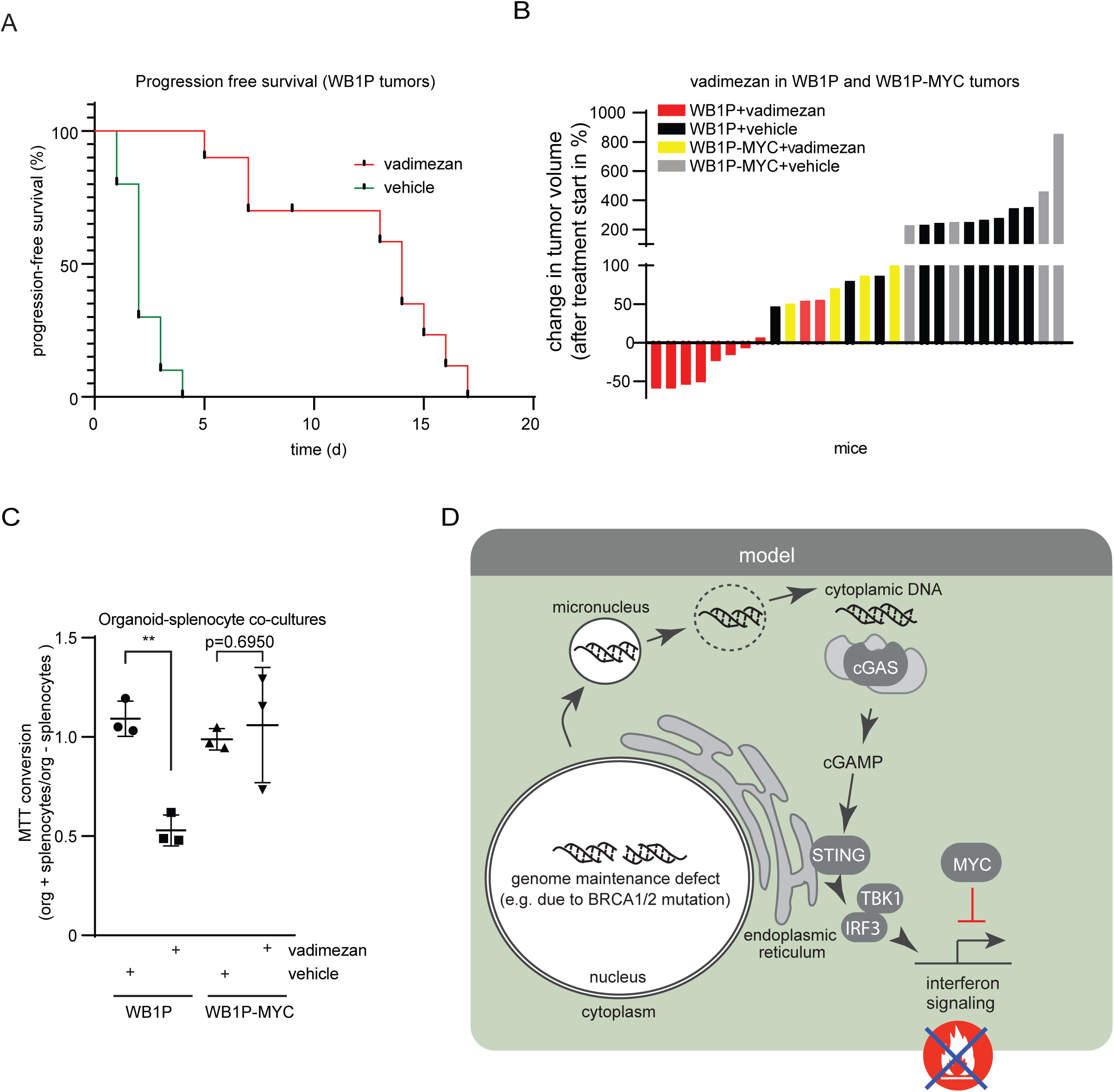
MYC suppresses anti-cancer efficacy of pharmacological STING activation in WB1P tumors. (A) Kaplan-Meyer-curves of mice transplanted with syngeneic WB1P tumor organoids and treated once with vehicle or 25mg/kg vadimezan i.p when tumors reached 100mm^3^. Shown is the time of progression free survival after treatment. (n=10) (B) A waterfall plot depicting changes in tumor volumes 7 days after treatment as described in A. Mice transplanted with syngeneic WB1P (n=10) as well as WB1P-MYC (n=4) organoids are shown. (C) Organoid-splenocyte co-cultures were checked for viability with an MTT-assay with and without vadimezan treatment (10ug/ml, 7 days culture, n=3). Organoid viability as measured by color intensity from the MTT assay was used as read-out. Plotted is the ration of color intensity of organoids cultured with spenocytes versus organoids without them, demonstrating that MYC expression protects organoids from splenocyte attack during vadimezan treatments (students t-test, p=0.0012, n=3 independent replicates, average of 5 wells/condition/replicate is plotted). (D) Model illustrating how MYC suppresses STING/IFN signaling in tumor cells with genomic instability.

## Discussion

It remains enigmatic why TNBCs and *BRCA*-mutated breast cancers rarely respond to immunotherapy (Voorwerk et al., 2019) despite being viewed as immunogenic due to the increased amount of neo-antigens induced by their genomic instability (Turner et al., 2004), (Solinas et al., 2019), (Woo et al., 2015), (Kraya et al., 2019). In this study, we show that a substantial fraction of TNBCs may evade immune clearance via MYC overexpression. We show that MYC suppresses immune cell infiltration into tumors by repressing IFN/STING signaling in a tumor cell-intrinsic manner. Our findings provide one mechanism by which TNBCs and *BRCA*-mutated breast cancers may evade clearance by the immune system.

Suppression of immune responses against tumor cells is of great importance for cancer development and progression. Consequently, therapeutic boosting of the adaptive immune system via inhibition of immune checkpoint components PD1/PD-L1 and CTLA4 has been successfully used to activate T cell responses against tumors (Ribas and Wolchok, 2018). Surprisingly, the anti-PD-1 inhibitor pembrolizumab was effective in only 20% of TNBC patients as monotherapy (Voorwerk et al., 2019), despite the fact that TNBCs are predicted to be immunogenic due to high levels of genomic instability (Nanda et al., 2016), (Adams et al., 2019). Also, no association was found between response to pembrolizumab and *BRCA1/2* mutations, and BRCA1-like genomic copy number profiles were even negatively associated with response (Voorwerk et al., 2019). This is especially puzzling since loss of BRCA1/2 function results in activation of STING signaling and subsequent attraction of immune cells in patients (Heijink et al., 2019; Parkes et al., 2017; Reisländer et al., 2019). Our results suggest that the efficacy of immune checkpoint inhibitor (ICI) therapy in TNBC and *BRCA*-mutated breast cancer may be restrained by MYC-induced suppression of local immune cells. This may be the case in a substantial fraction of TNBCs, as our analysis of oncogene amplification in a cohort of 142 human *BRCA*-mutant TNBCs from the TCGA dataset revealed *MYC* amplifications in the majority of cases, far more often than any other oncogene. GSEA of RNA-seq data from the same cohort showed association of *MYC* amplification with reduction in immune signatures, specifically IFN and inflammatory signaling. Complementary GSEA of a cohort of samples from the TONIC trial (Voorwerk et al., 2019) confirmed the correlation of high MYC expression with reduced immune signatures, in line with previous findings, where *MYC* amplifications were correlated with immune deserts in TNBCs (Xiao et al., 2019).

Previous studies have shown that the effects of MYC on the tumor immune microenvironment differ between cancer types. For example, MYC activation induces CCL9-mediated attraction of macrophages in both lung adenocarcinoma (LUAD) and pancreatic ductal adenocarcinoma (PDAC). Still, it has opposite effects on B cells in these cancer types, driving their expulsion in LUAD and influx in PDAC (Kortlever et al., 2017), (Sodir et al., 2020). Another study reported that MYC activation in PDAC represses IFN signaling and invasion of NK and B cells via transcriptional repression of IRF5/7 and STAT1/2 (Muthalagu et al., 2020). Together, these studies suggest that MYC has tissue-specific effects on distinct immune cell populations via defined signaling pathways. Our study shows that in triple-negative and *BRCA1/2*-mutated breast cancer, MYC utilizes yet another strategy to promote immune evasion, namely by expulsion of virtually all tumor-infiltrating immune cell populations. Our data also show that MYC does not act via a single downstream target, but instead functions as a master regulator, inhibiting numerous effectors of multiple immune-signaling cascades.

To enhance the efficacy of ICI therapy, an obvious goal is turning immunologically “cold” tumors into “hot” tumors (Li et al., 2016), (Iurescia et al., 2018). A potential strategy to bypass the immune-suppressive effects of MYC overexpression in triple-negative and *BRCA*-mutated breast cancer would be to activate IFN signaling via STING agonists in tumor cells and their environment (Ramanjulu et al., 2018), (Weiss et al., 2017). However, while boosting STING signaling inhibits growth of TNBCs that do not overexpress MYC, its impact on MYC overexpressing TNBCs is rather moderate, supporting our notion that MYC targets effectors downstream of STING and effectors of other immune pathways. Effective inhibition of the immune-suppressive effects of MYC overexpression in TNBC would therefore require activating innate immunity at a more downstream level via administering interferons or direct targeting of MYC activity, which has proven challenging thus far.

It has recently been reported that not only MYC, but also other oncogenes such as KRAS may have the ability to suppress immune responses against tumors by inhibiting IFN signaling (Liao et al., 2019). In line with this, we find that amplifying mutations in several oncogenes in human TNBC leads to markedly decreased expression of immune signature genes, including genes related to IFN signaling.

Taken together, our findings demonstrate a role for MYC in counteracting immune-cell invasion in TNBC via direct inhibition of IFN signaling responses. MYC-induced expulsion of TILs could explain the ineffectiveness of ICI therapy in a large fraction of triple-negative and *BRCA*-mutated breast cancers that are potentially immunogenic due to their genomic instability. Insight into how IFN signaling is silenced should be incorporated into designing combination therapies to activate IFN signaling and ultimately improve the response rates of MYC-amplified TNBCs to ICIs.

## Acknowledgments

We are grateful for excellent support from the NKI animal facility, RHPC computing facility, flow cytometry facility, animal pathology facility, transgenic facility, preclinical intervention unit, core facility molecular pathology and biobanking (CFMPB), and genomics core facility. We want to thank Martine van Miltenburg for providing data regarding the KP tumor model and Ivo Huijbers for help with generating the mouse models. This work was carried out on the Dutch national e-infrastructure with the support of SURF Cooperative (e-infra160136). Financial support was provided by the Oncode Institute, the Netherlands Organization for Scientific Research (NWO: Cancer Genomics Netherlands (CGCNL), VICI 91814643 (J.J.), VICI 91819616 (K.E.dV.), VIDI 91713334 (M.A.T.M.v.V.)), the European Research Council (ERC Synergy project CombatCancer (J.J.), ERC-Consolidator grant “TENSION” to (M.A.T.M.v.V.), ERC-Consolidator award InflaMet 615300 (K.E.dV.)), a National Roadmap grant for Large-Scale Research Facilities from NWO (J.J), the Dutch Cancer Society KWF10083 and KWF10623 (K.E.dV.), an unrestricted grant of the Hanarth Fonds (R.S.N.F) and an early postdoc mobility grant from the Swiss National Science foundation (D.Z).

## Author contributions

D.Z: conceptualization, investigation, data analysis, writing of original draft, review and editing C.B: conceptualization, investigation, data analysis, writing of original draft, review and editing F.T: conceptualization, investigation, data analysis, writing of original draft, review and editing J.B: Data analysis

A.B: Data analysis

S.J: Methodology, data analysis A.M: Methodology, data analysis M.D.W: Methodology, data analysis K.K: Methodology, data analysis M.R: Data analysis

L.H: Methodology, data analysis R.dB: Data analysis

S.A: Methodology

E.vdB: Methodology AP.D.: Methodology C.L: Methodology M.vdW: Supervision L.W: Supervision K.E.dV: Supervision W.Z: Supervision

R.S.N.F: Supervision, review & editing, Methodology, data analyses, and Conceptualization M.A.T.M.vV: Conceptualization, supervision, writing of original draft, review & editing J.J: Conceptualization, supervision, writing of original draft, review & editing

## Materials and Methods

### Mice and *in vivo* procedures

*WapCre;Trp53^F/F^* (WP), *WapCre;Brca1^F/F^;Trp53^F/F^* (WB1P), *WapCre;Trp53^F/F^;Col1a1^invCAG-Myc-IRES-Luc/+^* (WP-Myc), *WapCre;Brca1^F/F^;Trp53^F/F^;Col1a1^invCAG-Myc-IRES-Luc/+^* (WB1P-Myc), *WapCre;Brca1^F/F^;Trp53^F/F^;Col1a1^invCAG-Met-IRES-Luc/+^* (WB1P-Met), and *WapCre;Brca1^F/F^;Trp53^F/F^;Col1a1^invCAG-Cas9-IRES-Luc/+^* (WB1P-Cas9) mice were generated as described previously (Annunziato et al., 2019a). *WapCre;Brca1^F/F^;Trp53^F/F^;Col1a1^invCAG-MycERT2-IRES-Luc/+^* (WB1P-Myc^ERT2^) were generated as described in (Annunziato et al., 2019a) for WB1P-Myc, the ERT2 was added in frame to the murine *Myc* sequence. Intraductal injections were performed as described (Annunziato et al., 2019a). In brief, lentiviral particles were injected intraductally into the mammary glands via the nipple of the mouse. After injection, mice were monitored for mammary tumors twice per week and sacrificed upon reaching humane end-points or tumor size of 1500mm3. Organoid transplantations into the fat pad of the 4^th^ mammary gland of syngeneic mice were performed as described (Annunziato et al., 2019a). For activation of Myc^ERT2^, Tamoxifen 400-citrate pellets were used as staple chow (Envigo, TD55125). Vadimezan was dissolved in 2.5% bicarbonate at 1mg/ml and administered at 25mg/kg once when tumors reached 100mm^3^. Animals were assigned randomly to treatment groups and the treatments were performed by animal technicians blinded regarding the hypothesis of the treatment outcome. All animal experiments were approved by the Animal Ethics Committee of the Netherlands Cancer Institute (Amsterdam, the Netherlands) and performed in accordance with the Dutch Act on Animal Experimentation.

### Human cell lines

Human breast cancer cell lines MDA-MB-231, HCC1806, BT-549 and HCC38 were obtained from ATCC (CRM-HTB-26, CRL-2335, HTB-122, CRL-2314). Breast cancer cell lines were cultured in Roswell Park Memorial Institute (RPMI) medium supplemented with 10% fetal calf serum and penicillin/streptomycin (100 units per mL). Human cell lines were cultured at 37 °C in a humidified incubator with 5% CO_2_.

### Viral vectors and transduction

The following vectors were already described in (Annunziato et al., 2019a). In brief, for Lenti-Cre-transduction, pBOB-CAG-iCRE-SD (Addgene, plasmid #12336) was used. Lenti-MycP2ACre and Lenti-Myc^ERT2^P2ACre were cloned as follows: GFP-T2A-puro was removed by AgeI and SalI digest from the SIN.LV.SF-GFP-T2A-puro (Kas et al., 2017) and P2ACre was inserted as AgeI-SalI fragment into the SIN.LV.SF-GFP-T2A-puro backbone. The murine *Myc* cDNA was isolated with BamHI-AgeI overhangs using standard PCR from cDNA Clone 8861953 (Source BioScience) and inserted into the SIN.LV.SF-P2ACre vector. Lenti-Myc^ERT2^P2A-puro was cloned by inserting the Myc^ERT2^ cassette via BamHI and Age1 into the SIN.LV.SF-GFP-T2A-puro backbone. The Lenti-sgPten, Lenti-sgNT, Lenti-sgPten-Myc and Lenti-sgNT-Myc vectors were generated by inserting the *Myc* cDNA with XbaI-XhoI overhangs into the pGIN lentiviral vector for sgRNA overexpression (Annunziato et al., 2019b). The non-targeting sgRNA (TGATTGGGGGTCGTTCGCCA) and sgRNA targeting mouse *Pten* exon 7 (CCTCAGCCATTGCCTGTGTG) were cloned as described (Sanjana et al., 2014). Sanger sequencing was used for validation of all vectors. Co-transfection of four plasmids was used to produce concentrated VSV-G pseudotyped lentivirus in 293T-cells (Follenzi et al., 2000). The qPCR lentivirus titration kit from Abm (LV900) was used to determine titers.

To generate dox-inducible knockdown cell lines, BT-549 and HCC38 cell lines were infected with Tet-pLKO-puro harboring short hairpin RNAs (shRNAs). Tet-pLKO-puro was a gift from Dmitri Wiederschain (Addgene plasmid #21915). Hairpin targeting sequences that were used are: BRCA1 (5′-GAGTATGCAAACAGCTATAAT-3′), BRCA2 (5′-AACAACAATTACGAACCAAACTT-3′), luciferase (‘shLUC’, 5′-AAGAGCTGTTTCTGAGGAGCC-3′). To generate MYC overexpressing cell lines, BT-549 and HCC38 cell lines were infected with retrovirus containing pWZL-Blast-myc. pWZL Blast myc was a gift from William Hahn (Addgene plasmid #10674). Lentiviral and retroviral particles were produced as described previously (Heijink et al., 2019). In brief, 293T packaging cells were transfected with 10 μg DNA in combination with the packaging plasmids VSV-G and ΔYPR or Gag-Pol and VSV-G complemented with pAdvantage using a standard calcium phosphate protocol. Virus-containing supernatants were harvested and filtered through a 0.45 μM syringe filter with 4 μg per mL polybrene. Supernatants were used to infect target cells in two or three consecutive 24-hour periods. Infected cells were selected in medium containing puromycin (2 μg per mL) or Blasticidin (1 μg per mL) for at least 48 hours. Monoclonal cell lines were grown after single-cell sorting. Knock-down or overexpression was confirmed by immunoblotting. For dox-inducible expression of MYC, cells were first transduced with pRetroX-Tet-On Advanced, and selected for7 days with 800 μg/mL geneticin (G418 Sulfate)(Thermo Fisher). Human CCNE1 was PCR amplified from Rc-CycE, which was a kind gift from dr. Bob Weinberg. Human MYC was PCR amplified from MSCV-Myc-T58A-puro, which was a kind gift from dr. Scott Lowe. CYCLIN E1 fragments were ligated into pJET1.2/blunt using the blunt end protocol from GeneJET (Thermo Fisher). MYC was digested with NotI and EcoRI and ligated into the corresponding cloning sites of pRetroX-Tight-Pur. Subsequently, cells were transduced with pRetroX-Tight-Pur harboring MYC and selected for two days with 5 μg/mL puromycin dihydrochloride (Sigma-Aldrich). To induce MYC expression, 1 μg/mL dox (Sigma-Aldrich) was added to the culture medium.

### Histology and immunohistochemistry

Tissues were formalin-fixed overnight and paraffin-embedded by routine procedures. Haematoxylin and eosin (HE) and immunohistochemical stainings were performed by standard protocols. The following primary rabbit antibodies were used for immunohistochemistry: anti-Myc (Abcam ab32072), anti-CD3 (Thermo Scientific, RM-9107), anti-F4/80 (abD serotec, MCA497) and anti-CD31 (AbCam ab28364). All slides were digitally processed using the Aperio ScanScope (Aperio, Vista, CA, USA) and captured using ImageScope software version 12.0.0 (Aperio).

### Generation of cGAS knockout cells

CRISPR guide RNAs were generated against cGAS (#1: 5’-caccgGGCATTCCGTGCGGAA-GCCT-3’; #2: 5’-caccgTGAAACGGATTCTTCTTTCG-3’) and cloned into the Cas9 plasmids pSpCas9(BB)−2A-Puro (PX459, Addgene #62988) and pSpCas9(BB)-2A-GFP (PX458, Addgene #48138) using the AgeI and EcoRI restriction sites. BT-549 and HCC38 cells were transfected with both plasmids simultaneously (2 μg) using FuGene (Promega) according to the manufacturer’s instructions. After transfection, cells were selected with puromycin (1 μg per mL) for 48 hours or single-cell sorted for GFP. Single-cell *CGAS^−/−^* clones were confirmed by immunoblotting. Subsequently, *CGAS^−/−^* or parental cells were infected with Tet-pLKO-puro shRNAs targeting BRCA1, BRCA2 or Luciferase as described before.

### Western blotting

Cultured cells were lysed in Mammalian Protein Extraction Reagent (MPER, Thermo Scientific), supplemented with protease inhibitor and phosphatase inhibitor cocktail (Thermo Scientific). Proteins were separated on SDS-polyacrylamide gels (SDS-PAGE) and transferred onto a polyvinylidene difluoride (PVDF) membrane (Millipore). Membranes were blocked in 5% milk or bovine serum albumin (BSA) in Tris-buffered saline, with 0.05% Tween-20. Immunodetection was done with antibodies directed against BRCA2 (1:1000, Calbiochem, #OP95), BRCA1 (1:1000, Cell Signaling, #9010), cGAS (1:1000, Cell Signaling, #15102), STING (1:1000, Cell Signaling, #13647), cMYC (1:200, Santa Cruz, sc40), pIRF3 (1:1000, Cell Signaling, # 29047), IRF3 (1:1000, Cell Signaling, # 4302), STAT1 (1:1000, Cell Signaling, # 9172), pSTAT1 (1:1000, Cell Signaling, # 8826) and beta-Actin (1:10.000, MP Biochemicals, #69100). Horseradish peroxidase-conjugated secondary antibodies (1:2500, DAKO) were used and visualized with chemiluminescence (Lumi-Light, Roche diagnostics) on a Bio-Rad Bioluminescence device equipped with Quantity One/Chemidoc XRS software (Bio-Rad).

### *In vitro* survival assays

BT-549 or HCC38 cells with indicated hairpins were plated in 6 wells (500 cells per well) and treated with or without dox (1 μg per mL) for 10-14 days. Cells were fixed in methanol and stained with 0.1% crystal violet in H2O. Plates were measured and quantified using an EliSpot reader (Alpha Diagnostics International) with vSpot Spectrum software. For proliferation assays, BT-549 and HCC38 cells with indicated hairpins were plated in 48 wells plates (10.000 cells per well) and cultured for up to 10 days with dox (1 μg per mL). At indicated time points, plates were centrifuged (900 RPM) for 10 minutes and cells were fixed with 10% Trichloroacetic acid (TCA) in H2O overnight at 4 degrees. Plates were washed with tap water and dried by air. Cells were stained with 0.1% Sulforhodamine B (SRB) 1% Acetic acid in H2O for 30 minutes at room temperature and subsequently washed with 1% Acetic acid-H2O. Bound SRB dye was dissolved by adding 10 mM Tris-H2O to wells and OD was measured at 510nM with an iMARK microplate reader (Bio-Rad).

### Quantitative RT-qPCR

Cell pellets from BT-549 and HCC38 treated with or without dox (1 μg per mL) for indicated time points were harvested and stored at -20 °C. RNA was isolated using the RNeasy Mini Kit (Qiagen) and complementary DNA (cDNA) was synthesized using SuperScript III (Invitrogen) according to the manufacturer’s instructions. Quantitative reverse transcription-PCR (RT-PCR) for cytokine mRNA expression levels was performed in triplicate using PowerUp™ SYBR™ Green Master Mix (Thermo Scientific). Primers used are listed in Table S1. Glyceraldehyde 3-phosphate dehydrogenase (GAPDH) was used as reference gene and experiments were performed on an Applied Biosystems Fast 7500 machine.

### ELISA

To analyze cytokines and chemokines secreted by breast cancer cells, BT-549 and HCC38 cells with indicated hairpins were treated with or without dox (1 μg per mL) and plated at similar cell densities. Cell culture media was harvested at indicated time points and stored at -20. According to the manufacturer’s instructions, concentrations of CCL5 (R&D Systems, DY278-05) were measured using Enzyme-Linked Immuno Sorbent Assay (ELISA) according to manufacturer’s instructions.

### Cytokine and chemokine array

According to the manufacturer’s protocol, proteome profiler Mouse XL Cytokine array (R&D system) was performed on whole cell lysate from WB1P and WB1P organoids, according to manufacturer’s protocol.

### Immunofluorescence microscopy

Cells were grown on coverslips and treated with or without dox (1 µg per mL) for indicated time points. For RAD51 foci formation, cells were irradiated with 5Gy using a CIS international/IBL 637 cesium137 source. After 3h of irradiation, cells were washed with PBS and fixed in 2% formaldehyde with 0.1% Triton X-100 in PBS for 30 min at room temperature. Cells were permeabilized in 0.5% Triton X-100 in PBS for 10 min and subsequently blocked with PBS containing 0.05% Tween-20 and 4% BSA for 1 h. For micronuclei staining, cells were fixed in 4% formaldehyde for 15 min at room temperature. Subsequently, cells were permeabilized with 0.1% Triton X-100 in PBS for 1 min followed by blocking in 0.05% Tween-20 and 2.5% BSA in PBS for 1 h. Cells were incubated overnight with primary antibodies against RAD51 (1:400, GeneTex, #gtx70230), Geminin (Cell Signaling, #9718, 1:200) or cGAS (1:200, Cell Signaling, #15102) in PBS–Tween–BSA. Cells were extensively washed and incubated for 1 h with Alexa-conjugated secondary antibodies (1:400) at room temperature in the dark. Slides were mounted with ProLong Diamond Antifade Mountant with DAPI (Thermo Scientific). Images were acquired on a Leica DM-6000RXA fluorescence microscope, equipped with Leica Application Suite software.

### ChIP-seq

Duplicate samples were used for Chip-seq data generation. Organoids were cultured in 15cm dishes. Medium was replaced by PBS containing 1% PFA and plates were left shaking for 10 min at RT.

Di(N-succinimidyl) glutarate (DSG) (2mM) was then added and left shaking for 25 min after which reactions were quenched with 2.5M glycine for 5 min. Organoids were then washed with ice-cold PBS + protease inhibitor (Roche). ChIP and sample processing was performed as described previously (Singh et al., 2019). Five μg of cMYC antibody Y69 (Abcam, ab32072) and 50 μl of magnetic protein A (10008D; Thermo Fisher Scientific) were used per IP. For ChIP-seq of tumor tissue, OCT-embedded tumors were cut in 30um sections and processed as described (Singh et al., 2019). The prepared libraries were sequenced with 65 base single reads on Illumina Hiseq 2500. The sequencing reads were aligned to the mouse genome GRCm38 (mm10) using Burrows-Wheeler Aligner (BWA, v0.7.5a; (Li and Durbin, 2009) with a mapping quality >20. Peak calling was performed using MACS2 v2.1.1.20160309 (q-value threshold 0.01, extension via Phantom Peaks). For each organoid and tumor dataset, the peaks from duplicate samples were merged based on the peak ranges using ChIPpeakAnno v3.18.2 (Zhu et al., 2010) and considered as MYC binding loci. The gene closest to each merged peak was defined as MYC target based on the GRCm38 (mm10) genome annotation. Data deposited under ENA accession number “PRJEB43214”.

### Flow cytometry

Tissues were collected in ice-cold PBS. Blood samples were collected in tubes containing heparin (Leo Pharma) and treated with red blood cell lysis buffer (155mM NH4CL, 12mM NaHCO3, 0.1mM EDTA) (RBC). Tumors were mechanically chopped using a McIlwain Tissue Chopper (Mickle Laboratory Engineering) and digested either for 1 hour at 37°C in a digestion mix of 3 mg/ml collagenase type A (Roche, 11088793001) and 25 μg/ml DNAse (Invitrogen, 18068–015) or for 30 min at 37°C in 100 μg/ml Liberase (Roche, 5401127001), in serum-free DMEM (Invitrogen). Reactions were terminated by addition of DMEM containing 8% FCS and cell suspensions were dispersed through a 70 μm cell strainer (BD Falcon, 352350). All single-cell suspensions were treated with RBC lysis buffer to remove red blood cells. Single-cell suspensions were plated in equal numbers in round bottom 96-wells plates (Thermo Scientific). Cells were incubated with mouse Fc Block^TM^ (BD Biosciences) for 15 min at 4°C and subsequently incubated with different combinations of fluorescently labeled monoclonal antibodies for 20 min in the dark at 4°C.7AAD viability staining solution (eBioscience, 00–6993) was added to exclude dead cells. Flow cytometric analysis was performed on a BD LSRII using Diva Software (BD Biosciences). Data analyses were performed using FlowJo Software version 10.0 (Tree Star Inc.).

The following antibody panels were used: Myeloid panel – CD45-eFluor605NC (1:100; clone 30-F11, eBiosciences), CD11b-eFluor650NC (1:400; clone M1/70, eBiosciences), Ly6G-AlexaFluor700 (1:200; clone 1A8; BD Pharmingen), Ly6C-eFluor450 (1:400; clone HK1.4, eBiosciences), F4/80-PE (1:200; clone BM8, eBiosciences), CD49d-FITC (1:400; clone R1–2, eBiosciences), CD3 PerCP Cy5.5, CD206-FITC (1:200; clone C068C2, eBiosciences), 7-AAD (biolegend, cat. 420403); Lymphoid panel – CD45-eFluor605NC (1:50; clone 30-F11, eBiosciences), CD11b-eFluor650NC (1:400; clone M1/70, eBiosciences), CD3-PE-Cy7 (1:200; clone 145–2C11, eBiosciences), CD4-APC-eFluor450 (1:200; clone GK1.5, eBioscience), CD8- PerCP-eFluor710 (1:400; clone 53–6.7, eBiosciences, CD49b-APC (1:400; clone DX5, eBiosciences), CD19-eFluor780 (1:200; clone eBio1D3) 7-AAD.

For flow cytometry of mouse organoids and human cell lines BT-549 and HCC38, cells with indicated hairpins were cultured for different time points with dox and harvested by trypsinization and fixed with Fix buffer I (BD bioscience) for 30 min. on ice. Cells were washed with 1% BSA-PBS and permeabilized with Perm Buffer III (BD bioscience) for 30 min. on ice. Samples were washed with 1% BSA-PBS and incubated (150.000 cells per sample) with pIRF3 primary antibody (1:100, Cell signaling, #29047, clone D601M) for 1 hour at 4 °C and subsequently stained with AlexaFluor 488-conjugated goat anti-rabbit secondary antibody (1:300) for 1 hour at RT. Samples were measured on the FACS Calibur (Becton Dickinson), and data were analyzed using FlowJo software.

### RNA sequencing

Cells harboring a BRCA2 hairpin with or without MYC overexpression were treated with dox (1 μg per mL) for 6 days for Supplementary Figure 5. Cells were harvested and frozen at −80 °C. RNA was isolated using RNeasy Mini Kit (Qiagen) including DNase treatment. RNA was sent to Genomescan (Leiden, the Netherlands) for polyA-enriched mRNA sequencing using Illumina NovaSeq6000. Quality control of RNA samples consisted of fluorescent determination of sample concentration and fragment analysis. Samples were sequenced with 150 base-pair (bp) paired-end reads and generated 20 million reads per sample. FastQC and Samtools Flagstat software assessed RNA sequencing quality control. At least 80% of the bases had a Q-score ≥30. At least two or three biological replicates were used per cell line.

For RNA sequencing of mouse tumors, RNA was isolated from tumor pieces with the Qiagen RNA isolation kit. The mRNA library was generated using Illumina TrueSeq Stranded mRNA Library Prep Kit and sequenced with 65 base single-end reads on Illumina Hiseq 2500. The sequencing reads were aligned to the mouse genome GRCm38 (mm10) using TopHat v2.1 (Kim et al., 2013) and the number of reads mapped to each gene was quantified using HTSeq (Anders et al., 2015). DESeq2 v1.22.2 was used for read count normalization (median ratio method) and differential expression analysis. Genes with adjusted FDR<0.05 (Benjamini-Hochberg procedure) and |fold-changes|>1.5 were defined as differentially expressed genes. Data deposited under ENA accession number “PRJEB43214”.

### Single-cell RNA sequencing

Single cell sequencing was performed as described previously (Macosko et al., 2015) (http://mccarrolllab.org/dropseq/).

### Trans-well T cell migration assay

BT-549 and HCC38 cells with indicated hairpins were plated in 24-well plates (20,000 cells per well) and treated with dox (1 μg per mL) for 4 or 5 days. Human peripheral blood mononuclear cells (PBMCs) were isolated from peripheral blood from healthy volunteers by Ficoll-Paque density centrifugation (Ficoll-Paque PLUS, GE Healthcare Life Sciences) and enriched for CD8^+^ T cells with the MagniSort™ Human CD8^+^ T cell Enrichment Kit (#8804-6812-74, Invitrogen) according to manufacturer’s instructions. Enriched CD8^+^ T cells (750,000 cells per transwell) were added on top of the filter membrane of a transwell insert (6.5 mm Transwell with 3.0 μm pore, Corning) and incubated for 24 or 48 hours, after which supernatant from the lower chamber was harvested to quantify migrated T cells by microscopy.

### T cell proliferation assay

BT-549 and HCC38 cells were plated in 6-well plates (20,000 per well) and treated with dox (1 μg per mL) for 5 days. T cells were harvested and enriched for CD8^+^ T cells as described for the T cell migration assay. Enriched CD8^+^ T cells were stained with CellTrace Violet (#C34557, ThermoFisher) according to manufacturer’s instructions and cultured in 96-well plates (100,000 cells per well) with 200 μL conditioned medium harvested from breast cancer cells pre-treated with dox for 5 days. To activate T cells, T cells were co-cultured with Human T-Activator CD3/CD28 dynabeads (#11131D, Thermofisher) in a bead to T cell ratio of 1:4 or 1:8. For every condition, 2 wells were cultured and combined for analysis. At day of analysis, T cells were pooled, harvested, measured on the FacsVerse (BD Biosciences) and analyzed with FlowJo software.

### Organoid-splenocyte co-culture

Organoids were derived from WB1P or WB1P-Myc mammary tumors as described (Duarte et al., 2017). WB1P organoids were transduced with a lenti-GFP and WB1P-Myc with a lenti-mCherry lentivirus. Splenocytes were derived from FVB mouse spleen, by dissociation on a 70uM cell strainer. For the co-culture, 200,000 splenocytes and 10 organoids were plated together in a 24- well plate with 50% RPMI medium, 50% ENR medium, supplemented with IL-2 (Prepotech, 300IU/ml). Live cell imaging was performed with a Zeiss AxioObserver Z1 microscope for 7 days. Organoids areas were quantified using Zen software.

For the MTT assays, roughly 1000 cells, disrupted by a fire-hardened glass pipette to approximately 5-10 cells/clump were seeded together with 20000 splenocytes in a 96 well plate, using the same culturing conditions as described above. Vadimezan (MedKoo, #201050) was added at 10ug/ml for 7 days, then 3-(4,5-Dimethylthiazol-2-yl)-2,5-Diphenyltetrazolium Bromide, (ThermoFisher, #M6494) was added for 3h, followed by cell lysis for 16 hours in SDS lysis buffer. Plates were analyzed on a TECAN infinite M-plex plate reader.

### TCGA data preprocessing and quality control

Genes with a robust average gene expression (Hodges Lehmann estimate) lower than 20, were removed from the analysis. Differences in gene expression due to differences in cancer types were adjusted for every cancer type separately by performing the following steps for each gene: (i) robust average gene expression was obtained using Hodges Lehmann estimator; (ii) robust standard deviation of gene expressions were obtained using Hall’s estimator; (iii) gene expression was normalized using the following formula: Adjusted gene expression = (gene expression – robust average)/robust standard deviation.

### Differential gene expression analysis

To investigate the differential gene expression in the context of amplification of oncogenes, we retrieved DNA copy number data from The Cancer Genome Atlas (TCGA). For each of the oncogenes, the respective copy number profiles were used to classify samples as either amplified (log_2_(segment mean copy number) > 0.3) or neutral (0.3 ≥ log_2_(segment mean copy number) ≥ - 0.3). After that, Welch t-test was performed to identify differentially expressed genes upon amplification of each oncogene. A metric defined by (-log10(p-value)*sign(t statistic)) for each Welch t-test was obtained. The above analysis was done separately on the following sets of samples from TCGA: (i) all breast cancer samples; (ii) TNBC samples; (iii) breast cancer samples with high (mutation assumed to have a disruptive impact on protein) or moderate (mutation possibly changing protein effectiveness) mutation in either *BRCA1* or *BRCA2*; (iv) TNBC samples with a high or moderate disruptive mutation in either *BRCA1* or *BRCA2*.

### Gene Set Enrichment Analysis (GSEA)

For GSEA of oncogene-expressing and control BT-549 or HCC38 cells, genes were ranked based on the –log P value between oncogene-expressing cells and control cells (pBABE-empty). Genes enriched in oncogene-expressing cells were positive and genes enriched in control cells were negative. For GSEA of BRCA2-depleted cells with or without MYC overexpression, genes were ranked based on the -log P-value between MYC overexpressing cells and control cells. Genes enriched in MYC-overexpressing cells were positive and genes in control cells were negative. Gene sets of the Hallmark collection (MSigDB) were loaded into GSEA and analyzed in both cell lines. For GSEA of BRCA2-depleted cells with or without MYC overexpression, only significantly downregulated genes (p < 0.05) in MYC overexpressing cells were loaded into GSEA software for both cell lines. GSEA was performed utilizing 3 gene set databases (Hallmark, Reactome & Gene Ontology Biological Processes) from the MSigDB.v5.2 (Liberzon et al., 2015). Gene sets containing less than 10 genes or more than 500 genes (after filtering out genes that were not present in our data sets) were excluded from further analysis. Enrichment of a gene set was tested according to the two-sample Welch’s t-test for unequal variance. Welch’s t-test was conducted between the set of metrics obtained from differential gene expression analysis of genes whose corresponding gene identifiers are members of the gene set under investigation and metrics of genes whose corresponding gene identifiers are not members of the gene set under investigation. To compare gene sets of different sizes, Welch’s t statistics were transformed to -log10(P-value). GSEA of mouse mammary tumors was performed based on the Wald statistic obtained from DESeq2 differential expression analysis using the fgsea Bioconductor package v1.8.0 (Korotkevich et al., 2016)). MsigDB Hallmark gene sets (Liberzon et al., 2015) with a minimum size of 15 and a maximum size of 3,000 were used for enrichment analysis.

### Differential immune cell type abundance

Immune cell type abundance in breast cancer samples from TCGA was estimated using CIBERSORT (Newman et al., 2019). The abundance of 22 immune cell types was estimated by applying the leukocyte gene signature matrix (LM22) on the mRNA expression profiles from TCGA. Raw values of the CIBERSORT analysis are listed in Supplementary Table 3. To investigate the differential immune cell type abundance in the context of amplification of MYC, we used DNA copy number data from TCGA to classify samples as either MYC-amplified (log_2_(segment mean copy number) > 0.3) or neutral (0.3 ≥ log_2_(segment mean copy number) ≥ - 0.3). After that, Welch t-test was performed to identify immune cell types that showed statistically significantly different abundance in MYC amplified versus neutral samples. A metric defined by (-log10(p-value)*sign(t statistic)) for each Welch t-test was obtained to explore the result. The above analysis was done separately on the following set of samples from TCGA: (i) all breast cancer samples; (ii) TNBC samples.

### Prediction of gene functionalities

A co-functionality network was generated with an integrative tool that predicts gene functions based on a guilt-by-association (GBA) strategy utilizing >106,000 expression profiles as described previously (Bhattacharya et al., 2020). The analyzer tool is available at http://www.genetica-network.com.

