## Supplementary Figure Legends for "MYC promotes immune-suppression in TNBC via inhibition of IFN signaling"

**Supplementary Figure S1 – Oncogene amplifications in human breast cancer**

**Related to Figure 1.**

(A) Frequency of amplified oncogenes in known *BRCA1/2*-mutated breast cancer samples (n=42) from TCGA data.
(B) Distribution plot of all breast cancer samples used for GSEA analysis from TCGA data. In total, 1028 breast cancer samples were included in the analyses. Individual samples are plotted on the x-axis, oncogenic aberrations on the y-axis.
(C) GSEA analysis of breast cancer samples (n=1082). Top 20 up (red) and down (blue) regulated hallmark genesets are plotted in amplified vs neutral samples for specific oncogenes. Number of amplified samples per oncogene is shown. Values plotted are Z-transformed p values.

**Supplementary Figure S2 – MYC activation alters tumor immunity in TNBC mouse models**

**Related to Figure 2.**

(A) Heatmap for the previously reported interferon-stimulated genes (Mackenzie et al., 2017) in RNA-seq of WB1P, WB1P-Myc, WP, and WP-Myc models.
(B) Heatmap for the previously reported MYC signatures (Bild et al., 2006) in RNA-seq of WB1P-Myc and WB1P bulk tumor, sorted tumor cells and organoids.
(C) Scatter plot of the correlation of expression changes between the WB1P and WP tumor models upon MYC overexpression. The correlation between MYC expressing tumors is independent of BRCA1 status.
(D) Flow cytometric analysis of CD3^+^, CD49b^+^, CD19^+^, CD11b^+^ presence in blood, spleen and lymph nodes of WB1P and WB1P-Myc mice showing no pronounced changes in peripheral immunity in the WB1P-Myc mice.
(E) Latency analysis of WB1P vs WB1P-Met vs WB1P-Myc tumors.
(F) Representative histogram of CD3 immunostaining in WB1P-Met tumor.

CD3+ cell counts are quantified for WB1P vs WB1P-Myc vs WB1P-Met tumors (see also Figure 2C).

**Supplementary Figure S3 – The shorter latency until tumor onset is not causal for the reduced immune infiltration in MYC overexpressing tumors**

**Related to Figure 2.**

(A) Heatmap of RNA-seq of WB1P-Myc vs WB1P-Met tumors. An IFN signature of 336 genes was used (Saleiro et al., 2015), showing the immunogenicity of WB1P-Met tumors.
(B) Representative images of CD3+ staining for tumors induced by intraductal injections of guides against PTEN with and without MYC overexpression. Right panels: Counts of CD3+ cells in defined areas.
(C) Scatter plot showing the correlation of expression changes between WB1P bulk tumors and sorted epithelial tumor cells by overexpression of MYC (R=0.83).
(D) Distribution of immune cell–type fractions in all breast cancer samples and TNBC only from TCGA were estimated with CIBERSORT analysis. Samples with amplified MYC (0.3 cut-off) were compared to samples with neutral MYC levels. Fractions of each immune cell type were compared with a Welch’s t-test. -log10(pvalue)*sign(t statistic) for each immune cell type are plotted. Color indicates a lower (blue) or higher (red) immune cell-type fraction in breast cancer samples with amplified MYC compared to neutral MYC samples.

**Supplementary Figure S4 – cGas/STING and IFN signaling is suppressed by MYC in mouse and human organoid and cell line models**

**Related to Figure 3.**

(A) Boxplots from RNA-seq data showing downregulation of CD74 and CIITA in WB1P and WB1P-Myc tumors.
(B) Western blot showing the absence of STAT1 expression in WB1P-Myc tumors.
(C) Cytokine array for CCL5 in WB1P and WB1P-Myc organoids.
(D) Representative image of flow cytometry analysis of p-IRF3 expression in WB1P-Myc^ERT2^ organoids with 1 day tamoxifen (dark grey), 5 days tamoxifen (light grey) and without (black), showing strong reduction of p-IRF3^+^ population upon MYC activation.
(E) Flow cytometry analysis of p-TBK1 in WB1P-Myc^ERT2^ organoids without tamoxifen (black), after 1 day with tamoxifen (dark grey) and after 5 days of tamoxifen treatment (light grey), showing strong reduction of p-TBK1^+^ population upon MYC activation.
(F) BT-549, BT-549-cGAS^-/-^, and HCC38 cells with indicated hairpins were depleted for BRCA1 or BRCA2, or overexpressed for MYC. Cells were treated with or without dox for three days prior to cell lysis, and immunoblotted for BRCA1, BRCA2, MYC, cGAS, STING and Actin.
(G) Representative image of long-term survival assay in BT-549. BT-549 cells harboring shBRCA2 with or without WZL-MYC were plated in 6-well plates and treated with or without dox. Cells were fixed after 10-14 days and stained with crystal violet.
(H) Quantification of long-term survival assay as described in G. BT-549 and HCC38 cells were plated in 6 wells with indicated hairpins with or without MYC overexpression and treated with dox. Cells were fixed and stained after 10-14 days. Percentage of cell survival was calculated by normalizing measurements to wells without dox treatment.
(I) Cell proliferation of BT-549 and HCC38 cells was analyzed with SRB assays. Equal numbers of cells were plated in 48 wells plates and treated with dox for several days. At indicated time points, cells were fixed and stained with SRB dye. OD values of dissolved SRB dye were normalized to OD value at day 0 of the same cell line.

**Supplementary Figure S5 – IFN signaling in TNBC human cell lines is suppressed by MYC**

**Related to Figure 3 and 4.**

(A) BT-549 and HCC38 cells with indicated hairpins, depleted for cGAS or overexpressed for MYC were treated with or without dox (1 μg per mL) for 6 days. RNA of cells was isolated and qRT-PCR was performed to analyze expression of IFN-γ, IFN-β1, CCL5 and CXCL10. GAPDH was used as reference gene. Fold changes were calculated with untreated conditions of each cell line. Mean fold changes are indicated underneath each condition. Error bars indicate SEM of at least three independent experiments with three technical replicates each.
(B) BT-549 cells with indicated shBRCA2 hairpin, depleted for cGAS or overexpressed for MYC were treated with or without dox for 5 days. Phosphorylation status of IRF3 was analyzed by immunoblotting.
(C) BT-549 shBRCA2 cells with or without MYC overexpression were treated for 5 days with dox and RNA-seq was performed. GSEA analysis with hallmark gene sets was performed on significantly downregulated genes in MYC overexpressed cells compared to BRCA2-depleted cells only. Means from three biological replicates per cell line were used for RNA-seq analysis. Enrichment scores of two examples are shown.
(D) HCC38 shBRCA2 cells were treated and analyzed as described in C. GSEA analysis with hallmark genesets was performed on significantly downregulated genes in MYC overexpressed cells compared to BRCA2-depleted cells only. Means from three biological replicates per cell line were used for RNA seq analysis. Enrichment scores of two examples are shown.
(E) Representative images of gating strategies from (un)activated T cells co-cultured with supernatant harvested from BT-549 or HCC38 cells. The activation of T cells was confirmed by their proliferation resulting in dilution of the violet celltrace marker per cell division as shown in the most right panel.

**Supplementary Figure S6 – Inducing MYC in existing tumors leads to expulsion of immune infiltrates**

**Related to Figure 4**

(A) Kaplan-Meyer curves of Myc^ERT2^-P2A-Cre injected B1P mice with and without tamoxifen.
(B) Relative tumor growth (left) of intraductally injected WB1P mice with Lenti-Cre MYC^ERT2^. Tamoxifen administration and immunostaining for CD3^+^ * (right), in selected mice with (*) showing that concomitant MYC de-activation and slower tumor progression is paired with increase of infiltrating lymphocytes and complete regression (CR) in one of the mice.
(C) Growth curves of WB1P-Myc^ERT2^ tumors with (black), without (green) and with tamoxifen from tumor size of 3x3 mm (red). On the right side are representative Micrographs of CD3 IHC in tumors with and without tamoxifen.
(D) WB1P organoids with a Myc^ERT2^ vector are orthotopically transplanted into mammary glands. Transplanted mice are treated with tamoxifen chow and resulting tumors assessed for CD3+ cell infiltration (IHC, CD3 in brown), quantification in right panel, n=5 mice/group, 5 windows/tumor counted, unpaired t-test, p=0.001

**Supplementary Figure S7 – Integration of ChIP-seq and RNA-seq of tumors and organoids shows MYC mediated direct inhibition of immunomodulatory factors downstream and in parallel of c-GAS/STING signaling**

**Related to Figure 5**

(A) Gene ontology analysis of differentially expressed genes in WB1P vs WB1P-Myc tumors, sorted epithelial tumor cells and organoids overlapped with MYC ChIP-seq peaks.
(B) Heat maps for the 59 downregulated genes involved in interferon signaling from the RNA-seq data of bulk, sorted tumor cells and organoids with a peak called in tumor and/or organoid ChIP-seq.
(C) Transcription factor motif enrichment of the sequences bound by MYC in the ChIP-seq in up-regulated, down-regulated and not transcriptionally affected genes, confirming MYC binding to its supposed target sequence.
(D) ChIP-seq reads in genomic region of *Irf9* and *Stat3*, two examples of genes with peaks in the promoter region.
(E) Constructed co-functionality network of genes upregulated by MYC (n=430) retrieved from overlapping MYC-ChIP-seq peaks with RNA sequencing data of WB1P and WB1P-Myc tumors and organoids. Genes share strong predicted co-functionality (r > 0.5) within network that was enriched with genes predicted to be involved in e.g. DNA repair and RNA processing.
