## Supplementary figures and images for "MYC promotes immune-suppression in TNBC via inhibition of IFN signaling"

### Supplementary Figure S1

Supplementary Figure S1

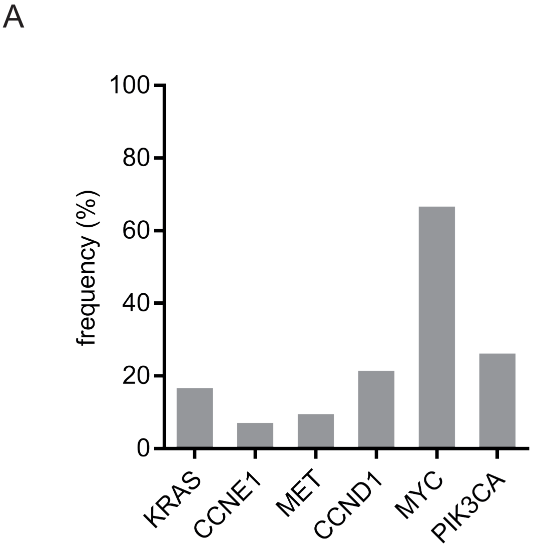


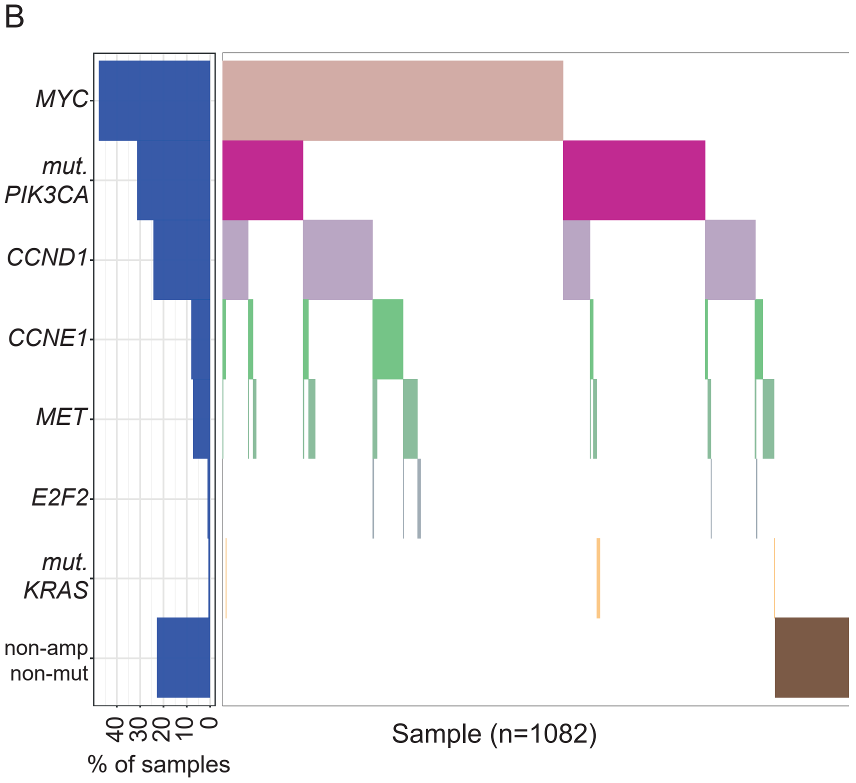


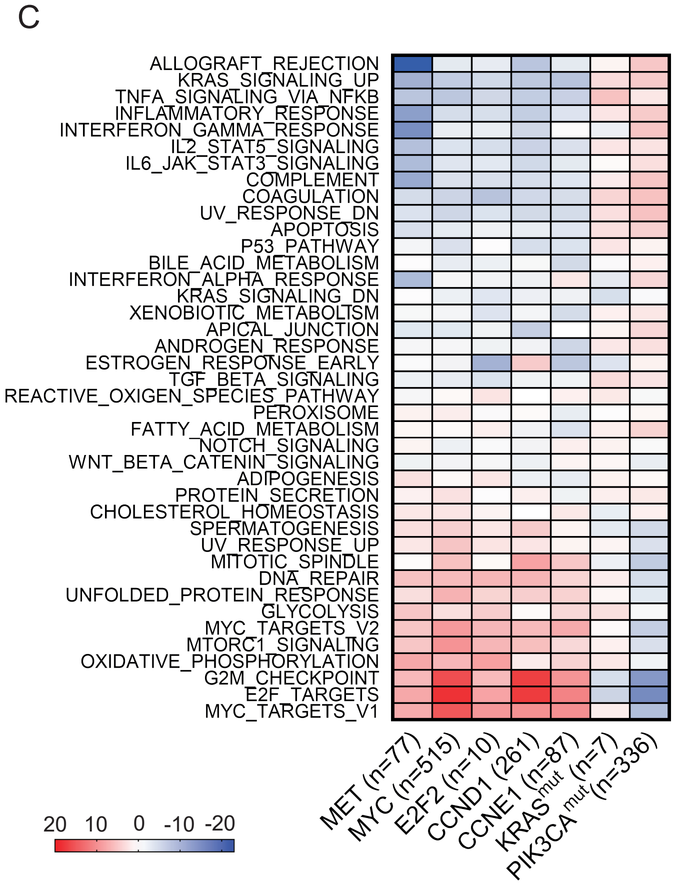

### Supplementary Figure S2

#### Supplementary Figure S2

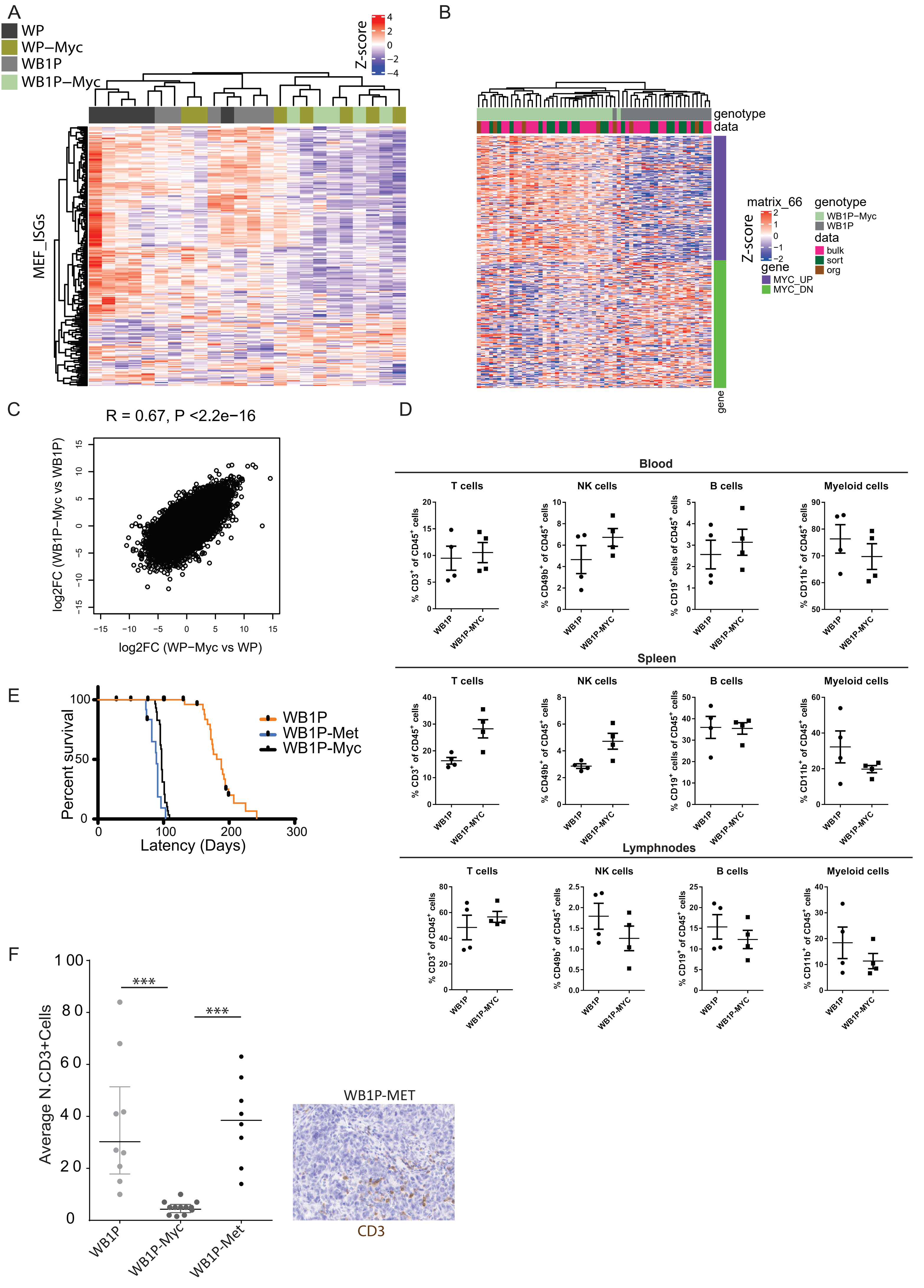

### Supplementary Figure S3

Supplementary Figure S3

A

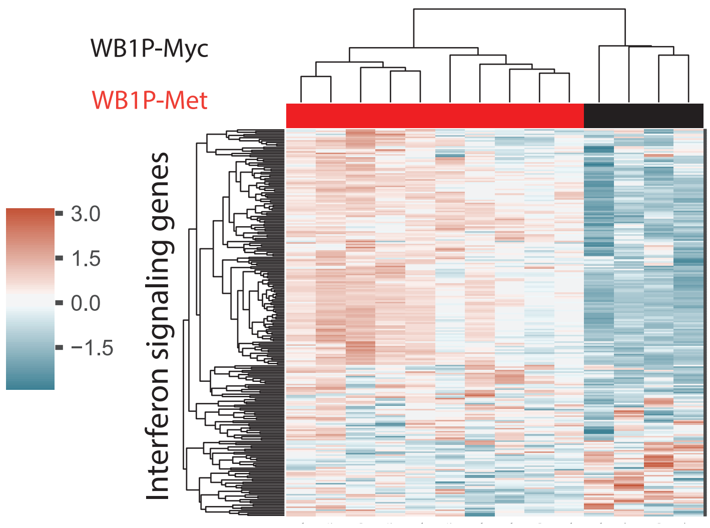


B

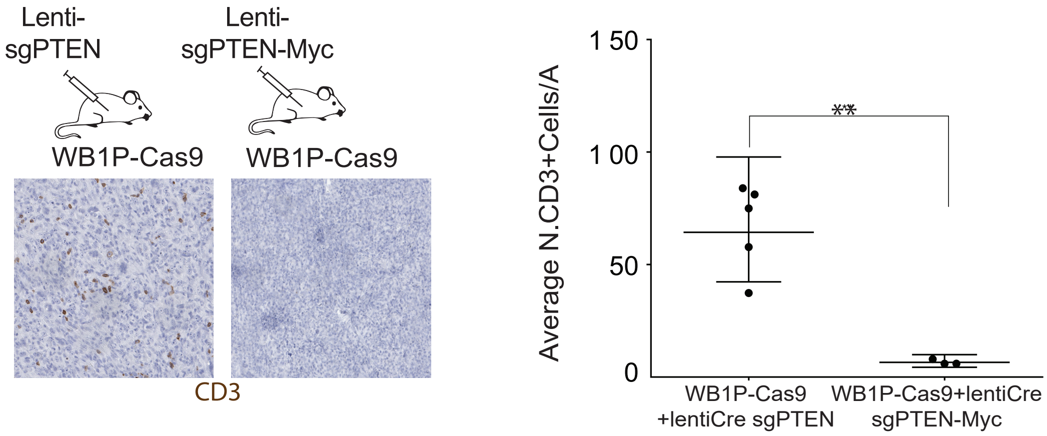


C

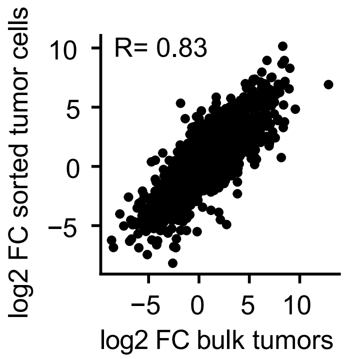


D

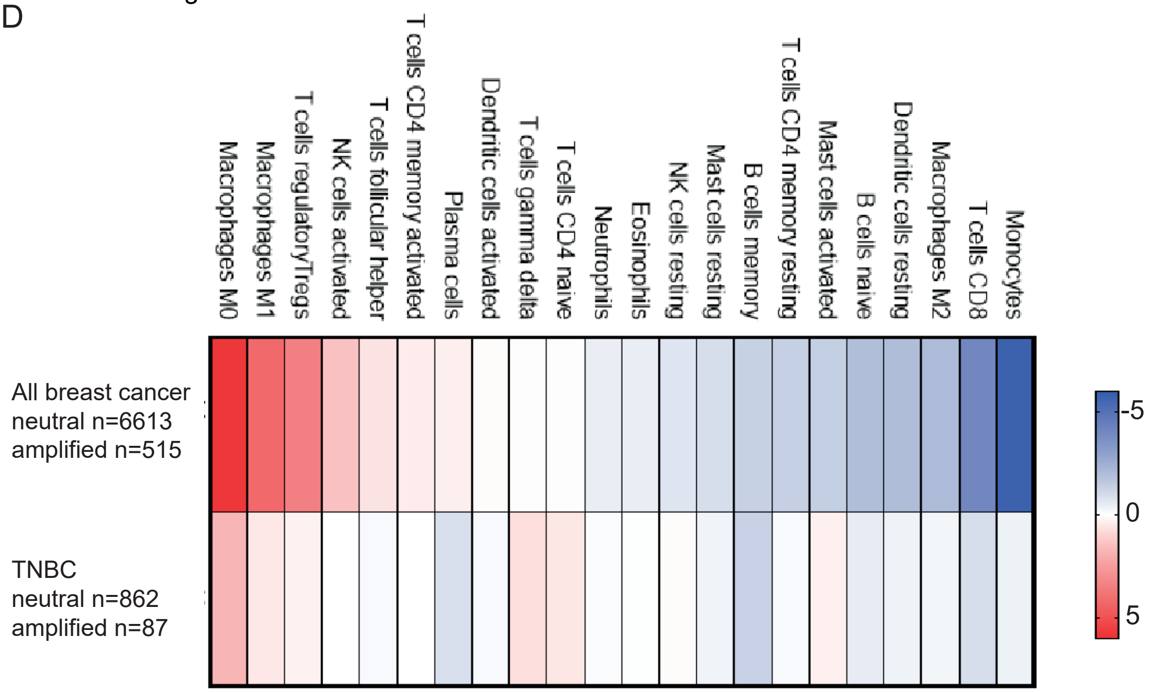

### Supplementary Figure S4

Supplementary Figure S4

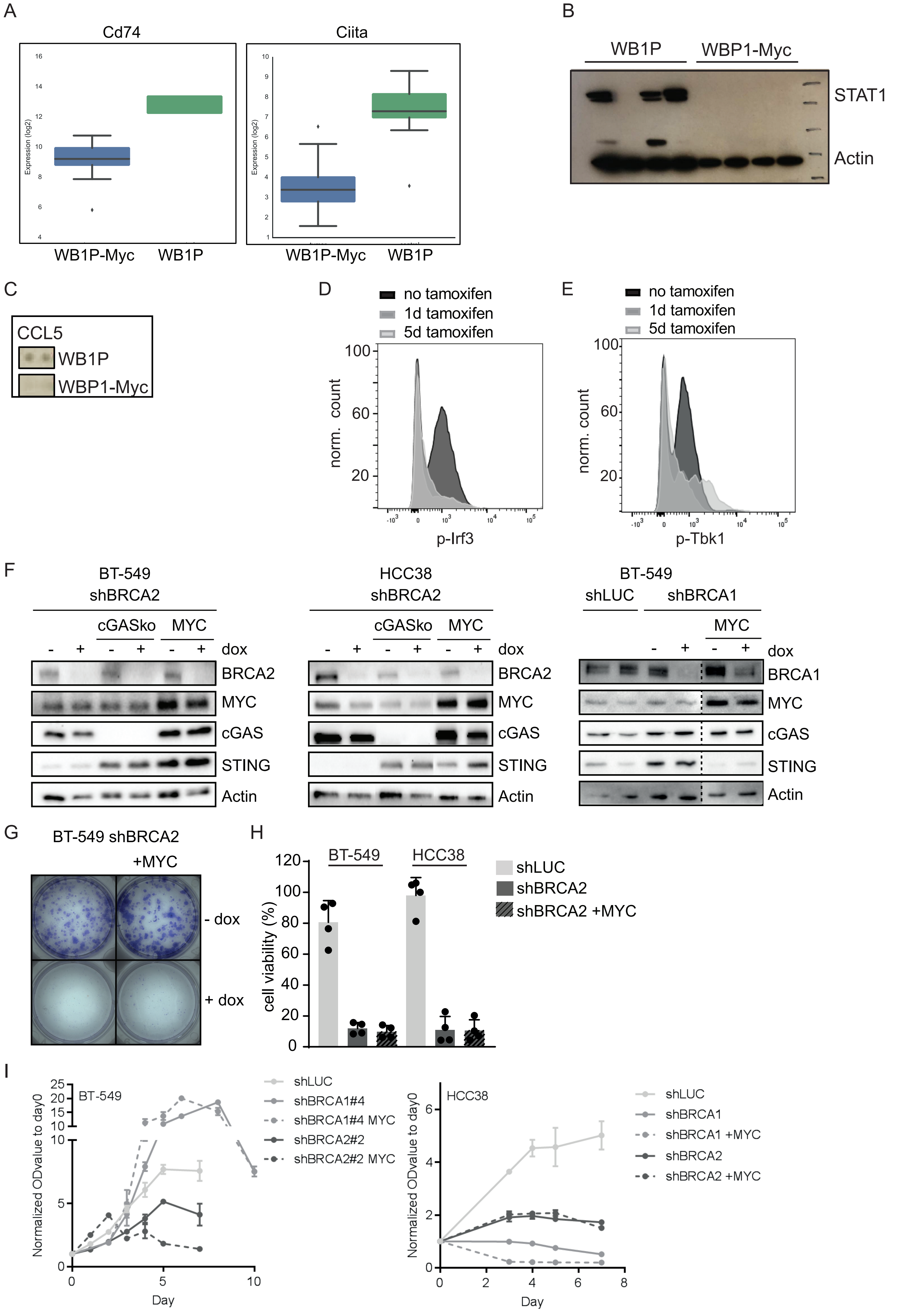

### Supplementary Figure S5

Supplementary Figure S5

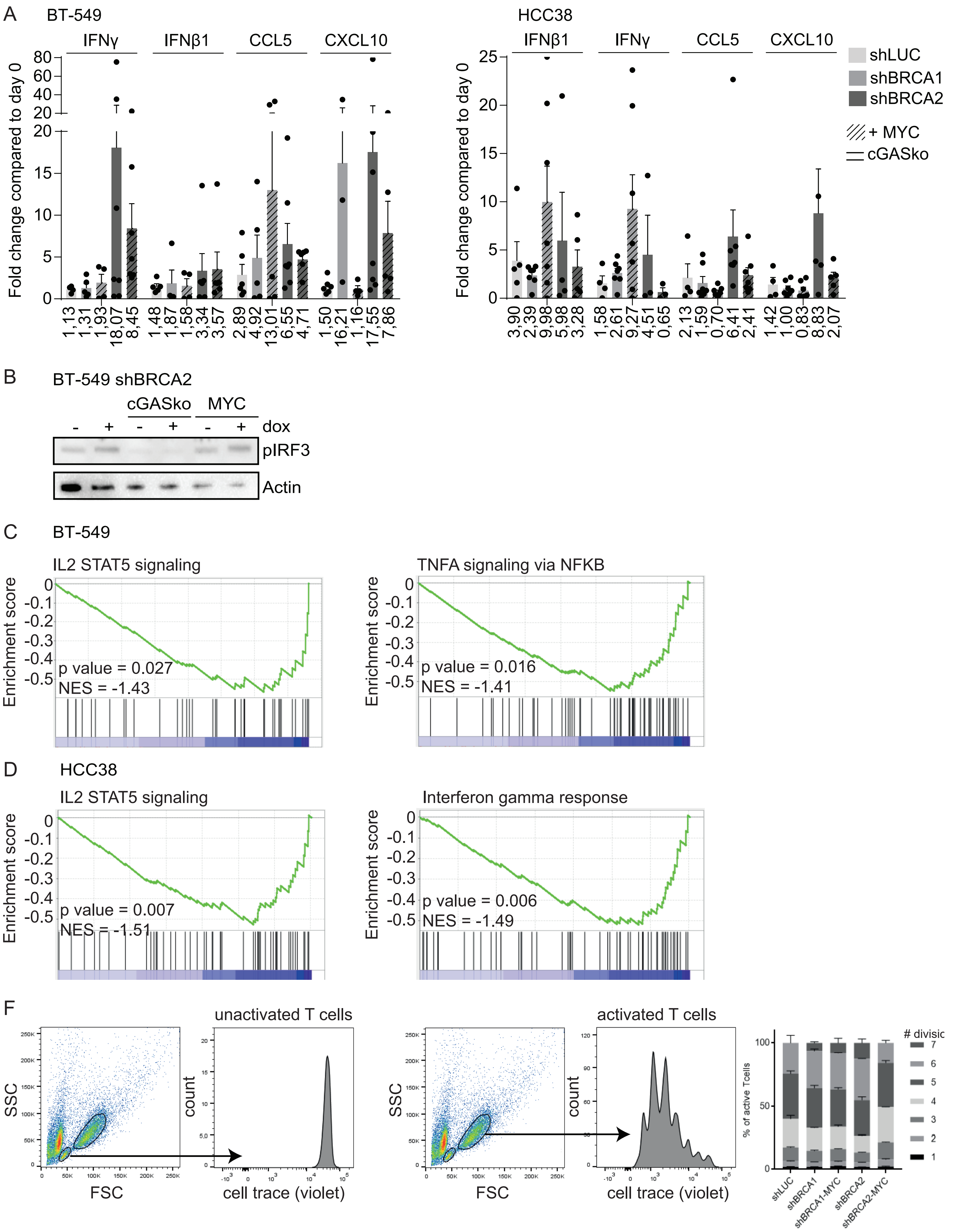

### Supplementary Figure S6

Supplementary Figure S6

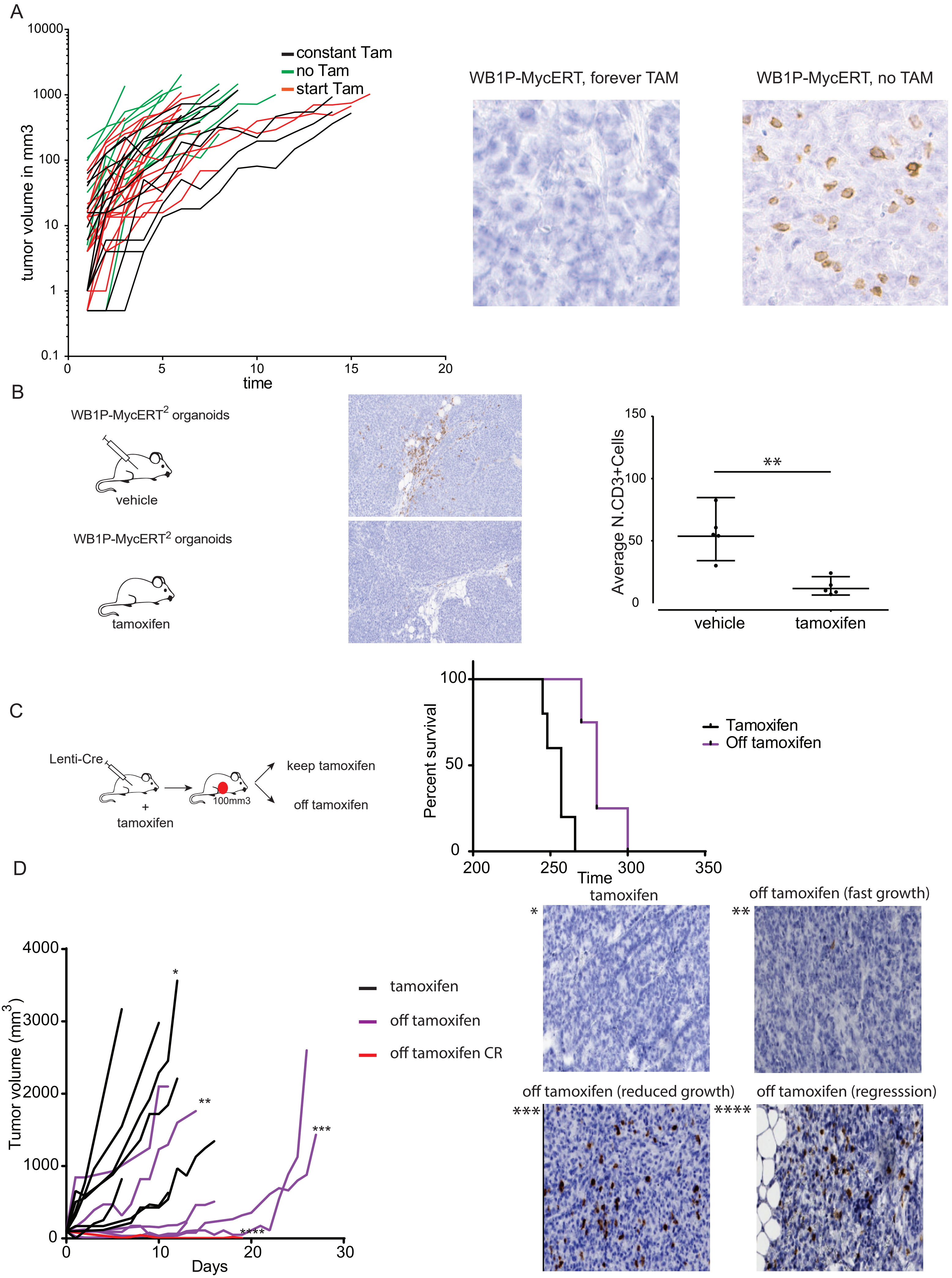

### Supplementary Figure S7

#### Supplementary Figure S7

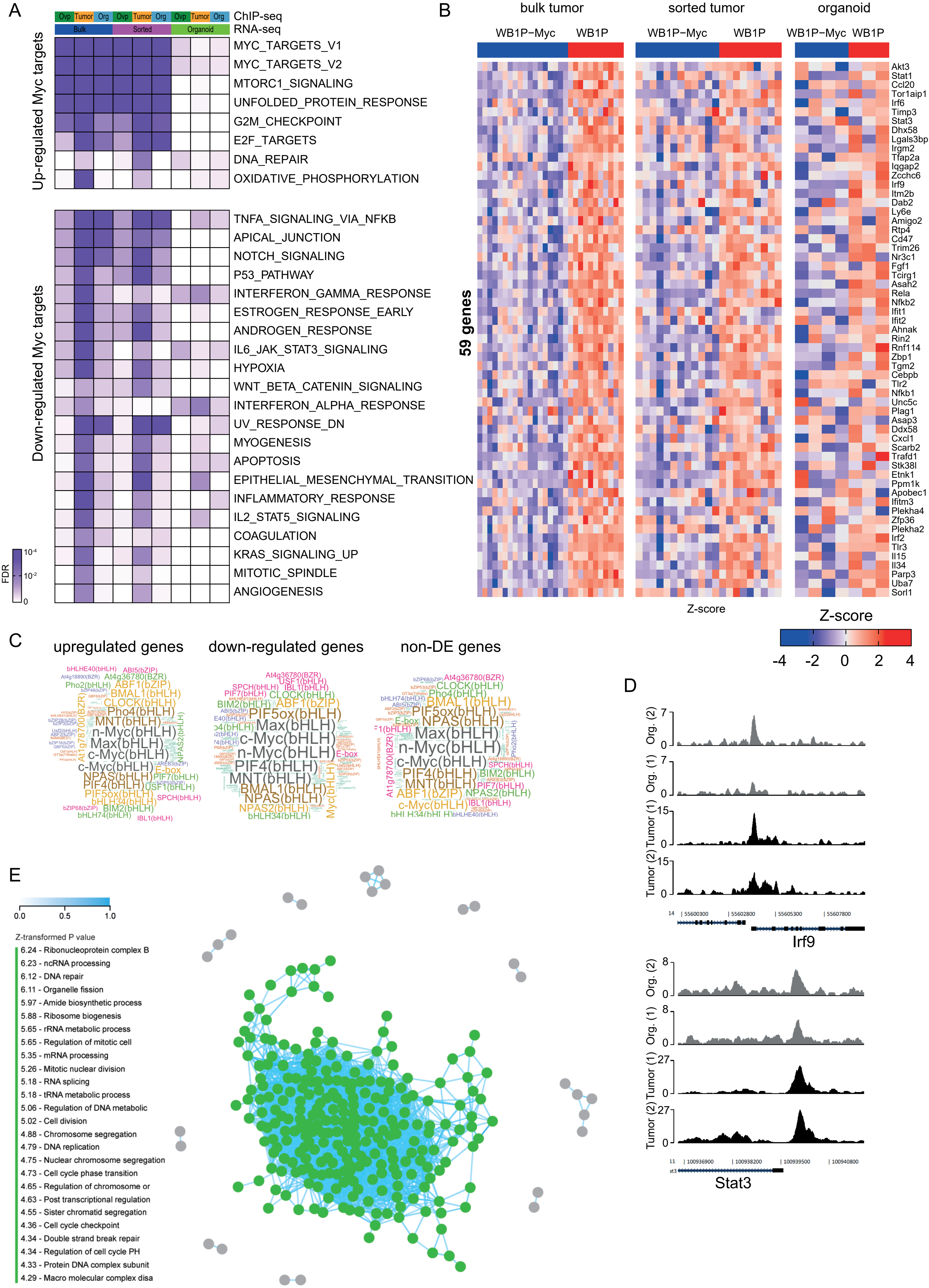
